# Cardiac Pacemaker Cells Harness Stochastic Resonance to Ensure Fail-Safe Operation at Low Rates Bordering on Sinus Arrest

**DOI:** 10.1101/2024.12.19.629452

**Authors:** Akihiro Okamura, Isabella K He, Alexander V Maltsev, Rostislav Bychkov, Syevda Tagirova, Michael Wang, Anna V Maltsev, Michael D Stern, Edward G Lakatta, Victor A Maltsev

**Affiliations:** National Institute on Aging, NIH, Baltimore, MD 21224, USA; School of Mathematics, Queen Mary University of London, London, UK

**Keywords:** Sinoatrial node, pacemaker, stochastic resonance, bradycardia, sinus arrest

## Abstract

**BACKGROUND:** The sinoatrial node (SAN) is the primary pacemaker of the heart. Recent high-resolution imaging showed that synchronized action potentials (APs) exiting the SAN emerge from heterogeneous signals, including subthreshold signals in non-firing (dormant) cells. This raises a new question in cardiac biology: how do these signals contribute to heartbeat generation? Here, we tested the hypothesis that pacemaker cells harness stochastic resonance to ensure fail-safe operation, especially at low rates bordering on sinus arrest.

**METHODS:** Membrane potential and Ca signals were measured using perforated-patch recordings in rabbit SAN cells exposed to sine-wave or white-noise currents. Additionally, we imaged Ca signals in intact mouse SAN tissue and performed multiscale model simulations at the subcellular, cellular, and tissue levels.

**RESULTS:** In addition to classical synchronized Ca transients, SAN tissue exhibited heterogeneous local Ca signals of different kinetics. Noise currents, mimicking the heterogeneous natural cell environment, restored AP firing in dormant cells and substantially improved the rate and rhythm of those firing infrequently and irregularly. The benefit followed a bell-shaped curve: the performance improved but then declined, demonstrating a hallmark of stochastic resonance. Rhythmic AP generation in response to sine-wave currents of different frequencies defined a resonance spectrum in SAN cells, reflecting their ability to respond via stochastic resonance to specific frequency components embedded in noise. Cholinergic stimulation shifted the resonance spectrum and responses to noise toward lower frequencies across all amplitudes tested, rendering cells unresponsive to higher-frequency signals while enabling more effective processing of slower signals. Both the numerical models and simultaneous recordings of membrane potential and Ca dynamics demonstrated that stochastic resonance is amplified by coupled electrical and Ca signaling, enhancing AP generation at low noise levels. Adding noise currents to the cell and tissue models allowed firing under conditions where they otherwise would have stopped.

**CONCLUSIONS:** SAN cells harness stochastic resonance amplified by coupled membrane-Ca signaling to ensure rhythmic heartbeat initiation, especially at low rates. This new signaling mechanism could help avoid sinus arrest when heart slows but noise increases, such as during parasympathetic stimulation, bradyarrhythmia, or aging.

## INTRODUCTION

The sinoatrial node (SAN) is a small region of heart tissue in the right atrium that generates rhythmic, spontaneous action potentials (APs) that pace cardiac contraction to meet the body’s demand for blood supply. Despite more than a century of research^1^, the origin of the heartbeat remains “still mysterious after all these years”^2^. This incomplete understanding of pacemaker mechanisms contributes, in part, to the fact that sinus node dysfunction, also known as sick sinus syndrome^3^, remains a major healthcare problem, especially in aged populations prone to bradyarrhythmia and life-threatening sinus arrest. Electronic pacemakers, currently used to treat sick sinus syndrome, impose lifestyle restrictions and can cause severe side effects.

Bradycardia occurs at the edge of normal SAN function and sinus arrest. What specific mechanisms maintain SAN function on this edge? The prevailing view of cardiac impulse initiation is that a small group of cells with the fastest AP firing rate then drives^4^ or entrains^5^ other SAN cells to deliver rhythmic impulses from the SAN exits. This view is consistent with earlier imaging of SAN tissue that showed concentric spread of excitation^6–8^. However, more recent imaging at single-cell resolution has revealed a new functional paradigm^9,10^. Excitation within the SAN center appears discontinuous and consists of functional cell communities (presumably signaling modules^11^) with distinct signaling patterns^9^. Some cells within the tissue generate no AP-induced transients^9,10^. Such non-firing cells (termed dormant cells^12^) represent a major population of cells enzymatically isolated from the SAN^12–15^. Dormant cells generate subthreshold noisy Ca and membrane potential oscillations, and many can be reversibly “awakened” to generate normal automaticity by β-adrenergic receptor stimulation^12,14–16^.

Thus, recent experimental data portray SAN pacemaking as an emergent property of a complex network of loosely connected, diversely firing and non-firing cells. The extreme structural heterogeneity of the SAN network, in which subthreshold irregular oscillations in some cells are mixed with AP firing in others, raises the possibility that random signal disturbances, often referred to as biological noise, could be an additional key factor in heart rate regulation^17,18^.

The role of biological noise in cardiac pacemaker function becomes even more apparent in light of experimental data suggesting that the heart’s pacemaker mimics brain cytoarchitecture and function^9,19,20^, and the cell cluster in the central SAN initiating the SAN impulse exhibits the most stochastic behavior^21^. Noise plays a fundamental role in information processing and affects all aspects of nervous-system function^22,23^. Because SAN tissue represents a complex network of interacting pacemaker cells and neuronal cells (the “little heart brain”^24^), it is reasonable to expect that in addition to channel noise within pacemaker cells and their membranes, noise sources found in neuronal networks, such as synaptic and sensory noise^22^, are also present in the SAN. For example, during cholinergic stimulation, individual local acetylcholine releases by nerve endings intensify and create disturbances in local signaling and AP-firing periodicity^25,26^. Additional noise sources influencing pacemaker cell function include ion-channel trafficking^27^ and connexin channels^28^.

Therefore, a frontier in cardiac biology is to understand the complex signal-processing mechanisms harbored within the heterogeneous SAN network that ultimately generate robust yet flexible cardiac impulses. Among those recently discussed^17,18,29–31^, one such signal-processing mechanism is stochastic resonance. Stochastic resonance is a phenomenon in which the addition of noise allows a subthreshold signal to become detectable by occasionally bringing the signal above threshold. Although stochastic resonance is recognized as a key information-processing mechanism in neuronal networks^32^, it has not been experimentally or theoretically demonstrated in SAN cells. Its presence and functional role in the SAN therefore remain unknown.

Each pacemaker cell exhibits intrinsic automaticity through diastolic depolarization generated by membrane currents coupled to periodic intracellular Ca signals within pacemaker cells^33^. Using combined experimental and theoretical approaches, the present study demonstrates that stochastic resonance is indeed a novel signal-processing mechanism in SAN cells that can support pacemaker function, especially at low AP firing rates both in basal state and during cholinergic receptor stimulation.

## METHODS

### Experimental methods and data availability

Detailed descriptions of experimental methods, materials, and statistical analyses, and numerical modeling are presented in the Supplemental Material. Experimental data associated with this study are available in Harvard Dataverse at https://doi.org/10.7910/DVN/PVEKM9.

## RESULTS

### Presence, patterns, and amplitude of Ca noise in intact SAN

To characterize the noise environment relevant to stochastic resonance in SAN tissue, we performed high-resolution, single-cell-level Ca imaging in intact mouse SAN preparations using two complementary methods: high-speed sCMOS camera (n=6 mice, Figure 1, Videos S1-S5), and confocal microscopy (n=3 mice, Video S6).

**Figure 1.**
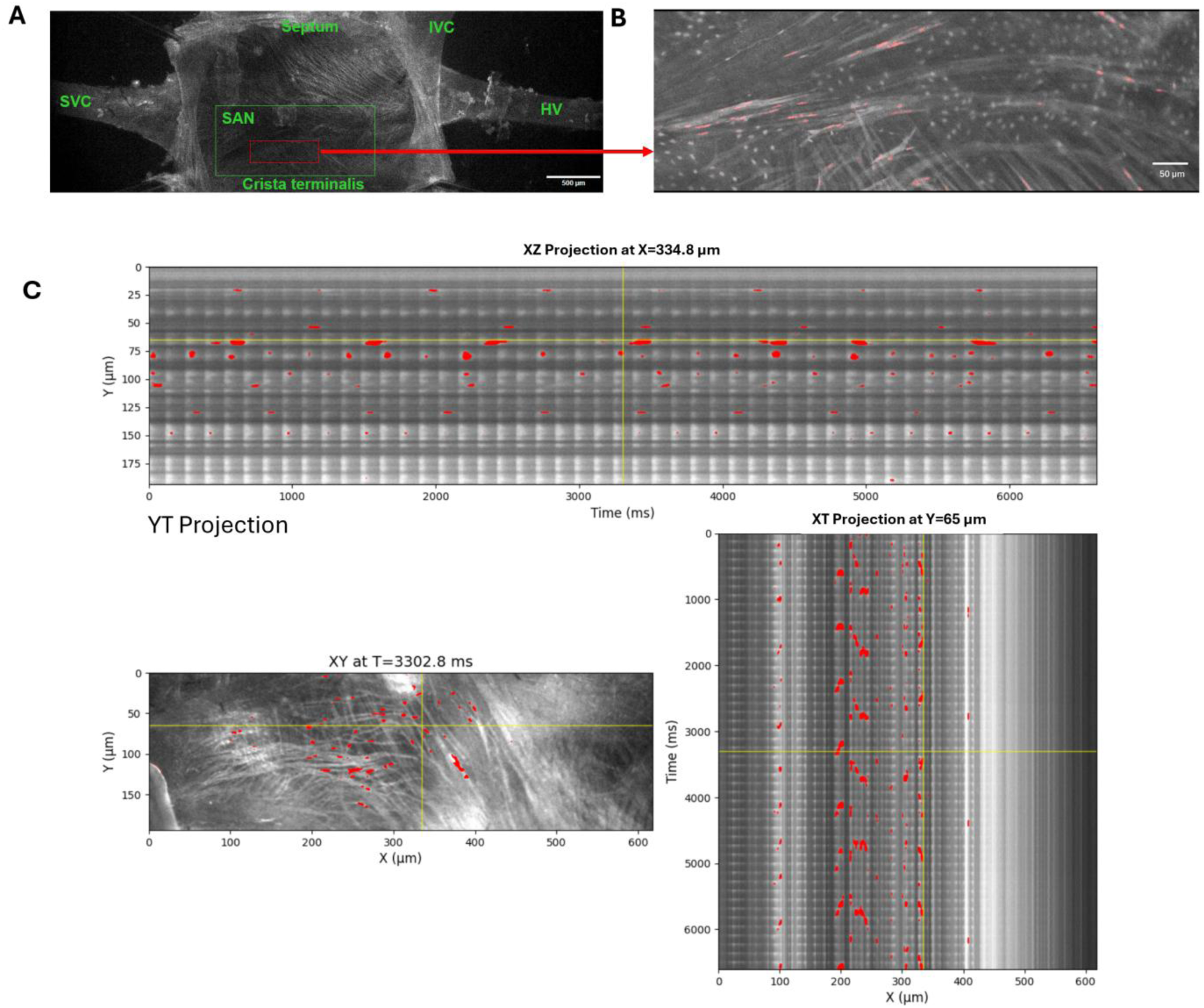
Presence of biological noise in intact SAN tissue. **A**, Low-zoom microscopic view of the entire mouse SAN preparation loaded with the Ca indicator Fluo-4, with anatomical benchmarks: crista terminalis, superior vena cava (SVC), hepatic vein (HV), and inferior vena ca**v**a (IVC). **B**, Higher-magnification view of a region of interest within the central SAN area. Scattered pink spots indicate background local Ca release (LCR) signals (Video S1). **C**, LCR signals (in red) detected in another SAN tissue preparation in a 2D image (XY) and in the corresponding XT and YT orthogonal projections (line-scan images along the yellow lines) to illustrate the spatiotemporal extent of the LCRs (Video S2). Our novel LCR-detection algorithm based on the PMD/Trefide/PCA signal-enhancement pipeline is described in detail in the Supplemental Material and Figure S1.

Figure 1 and Video S1 show the high-speed camera recordings of Ca signals in the SAN tissue at a low zoom with key anatomical landmarks. Imaging at higher resolution revealed that, in addition to the “global” AP-induced Ca transients (APCTs) visible as periodic synchronized flashes throughout the tissue (Video S1, upper panel), substantial heterogeneous local Ca signals were present between and during the APCTs. To objectively separate these background local signals from global transients, we developed a novel PCA-based image analysis pipeline (detailed in Supplemental Methods and Figure S1). Subtraction of the principal component (global APCT signal) from the total Ca activity revealed the residual biological noise generated by individual cells as local Ca release (LCR) signals (Figure 1B, Video S1 and S2 with LCRs highlighted in pink). Spatiotemporal dynamics of LCR signals different from rhythmic APCTs are also clearly seen in orthogonal projections of the Ca image stacks (line-scanning images, Figure 1C).

We identified three distinct patterns of background noise signals with different kinetics: 1) intracellular cell-width Ca waves propagating along cell length generated rhythmic, low-frequency local signals (Videos S1-S3); 2) multiple smaller and faster LCR signals occurred within individual cells (Video S4); 3) incoherent firing, i.e. local AP-like signals out of synchrony with the global transient (Video S5). The amplitudes of noise signals were substantial, comparable in magnitude to APCTs recorded simultaneously. The presence of all three patterns was confirmed by confocal microscopy, which provided higher spatial resolution within individual cells (Video S6).

### Functional AP firing range of SAN cells revealed by sine-wave protocols

Following isolation, each pacemaker cell manifests its own preferred AP firing frequency, and some can be dormant (i.e., non-firing; Figure S2). Here we examined the range of frequencies at which a cell can generate rhythmic APs when it is entrained (one-to-one capture) by external periodic signals. To this end, we tested responses of 63 SAN cells to a sequence of external currents in the form of sine waves 2*a*sin(2π*F*t) of various frequ*e*ncies (F) from 0.5 to 6 Hz and amplitudes (a; current amplitude in one direction) from 10 to 50 pA (Figure S3, top). The resonance spectrum of a given cell at a given perturbation amplitude was defined as the frequency range within which AP firing matched the frequency of the applied sine wave (criterion: ±0.1 Hz).

By applying a multitude of sine-wave protocols, we specifically measured the resonance spectra for four diverse cell populations: (i) fast-firing cells (>2.5 Hz, i.e. classical pacemakers); (ii) moderate-firing cells (1 to 2.5 Hz); (iii) slow-firing cells (< 1 Hz), and (iv) dormant cells firing no APs. Figure 2 shows examples of original recordings (panels A-D) and statistical analysis of the data (panels E-L). Our findings demonstrate that:

1. Larger-amplitude sine waves produced stronger effects, extending cell resonance spectra toward higher frequencies in a larger percentage of cells (blue bars, a = 50 pA, panels E-L).
2. All dormant cells began to fire APs as sine-wave amplitude increased.
3. The broadest resonance spectra were found for fast- and moderate-firing cells. The frequency range over which ≥50% of cells achieved one-to-one capture (dashed line in Figure 2, I-L) was the broadest in these cell populations (panels I and J) vs. that in slow-firing and dormant cells (panels K and L) at 25 and 50 pA amplitudes.
4. Inability of cells to follow low frequencies (0.5 and 1 Hz) signals led to burst firing: cells fired APs during the depolarizing part of the sine wave but fired no APs during the hyperpolarizing part (panels A-C). Conversely, dormant cells generated rhythmic firing at lower frequencies (as low as 0.5 Hz, panel D) but could not fire at higher frequencies (only 8% of cells could generate 5 Hz AP firing, panel L).
5. The resonance spectra shifted to lower frequencies as the intrinsic firing frequency of the cell population decreased. For example, at 50 pA amplitude the peak distribution was 3 Hz in fast-firing cells (blue bars, panel I), but only 2 Hz in slow-firing (blue bars, panel K).
6. Some dormant cells continued to fire after the sine-wave protocol ceased, displaying a “memory” effect (panel H, group “After”).
7. The resonance spectrum in each SAN cell is continuous. For example, cells maintained sine-wave capture as frequency increased in small 0.25-Hz steps, and capture occurred quickly, within 1 to 2 cycles (Figure S4).

**Figure 2.**
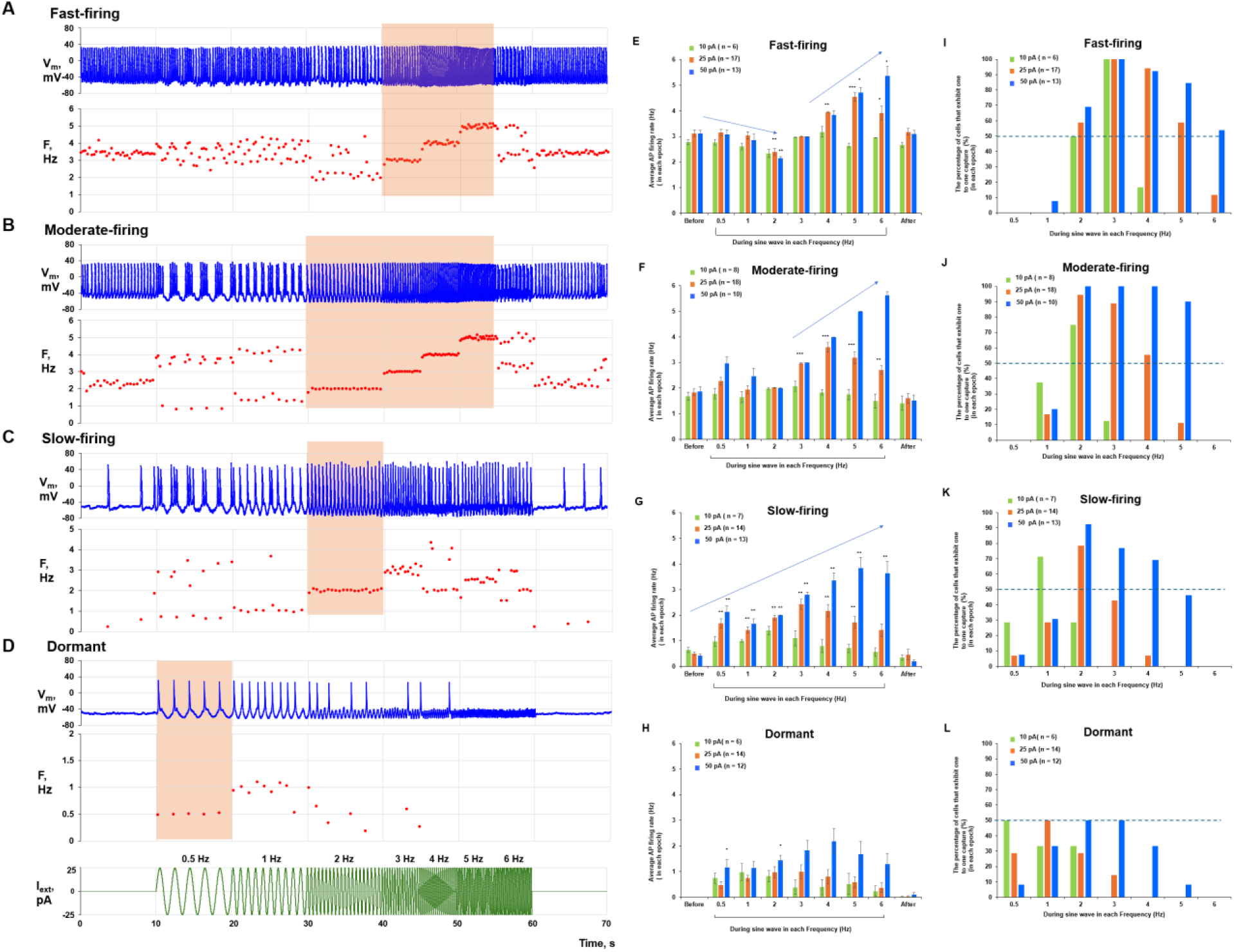
Resonance spectra of SAN cells revealed by sine-wave protocols. **A-D**, Representative examples of the effect of sine-wave currents (0.5 to 6 Hz, 25 pA; protocol shown at the bottom) on different cell populations. Original V_m_ traces and their corresponding intervalograms (in Hz) are shown. Resonance spectra are indicated by orange bands**. E-H**, Effects of sine-wave amplitude (color bars) on average AP firing rate (y-axis) at each sine-wave frequency (x-axis). Average AP firing rate in response to each sine-wave frequency was compared with that before sine-wave application using repeated-measures ANOVA or the Friedman test (*p < 0.05, **p < 0.01, ***p < 0.001). In addition, the Jonckheere-Terpstra trend test revealed significant trends in average rate change as wave frequency increased (arrows; tested separately and significant at p < 0.05 for 25 and 50 pA amplitudes). **I-L**, Percentage of cell**s** exhibiting one-to-one capture (y-axis) in response to each sine-wave frequency (x-axis) at each amplitude (colored bars). The horizontal dashed line indicates the 50% level and illustrates the stronger (broader) responsiveness of fast- and moderate-firing cells, especially at higher frequencies.

### Stochastic resonance in SAN cells spontaneously firing APs

The diverse noise signals (Videos S1-S6) of varying kinetics described above provide the biological substrate in SAN tissue for stochastic resonance, the subject of the study. This Ca release noise in the SAN likely occurs in concert with other potential noise sources mentioned in Introduction, such as neuronal mediator release^22,25,26^, ion channel trafficking^27^ and connexin channels^28^. White noise has a constant power spectral density (possessing equal intensity across frequencies) and therefore, in our study, serves as a representative model for the heterogeneous, noisy environment within the SAN. We tested the effect of white noise using four amplitudes: 25, 31.25, 62.5, and 125 pA. Because our protocol was zero-mean, current disturbances occurred with equal probability in both depolarizing and hyperpolarizing directions (Figure S3, bottom). Thus, the cell’s response was not inherently intuitive, given the complex nonlinear interplay of numerous ion channels and transporters with distinct Ca and voltage sensitivities and kinetics.

Panels A-C in Figure 3 show representative examples of the effect of noise on the firing frequency in each of 3 AP firing cell populations. Panels D-I show average AP firing frequencies and their coefficients of variation (defined as CV=SD/mean). An increase in CV indicates a decrease in rhythmicity and vice versa. Our findings are as follows:

1. For all populations of AP firing SAN cells and all tested noise amplitudes (except lower amplitudes of 25 and 31.25 pA in the fast-firing cells), the noise significantly increased the average AP firing frequency compared to intrinsic firing (orange bars vs. blue bars, Figure 3D-F). Furthermore, within each cell population, cells fired APs at significantly higher rates as the noise amplitude increased (statistical comparisons among orange bars in each cell population). Rarely, noise application decreased AP firing frequency at low noise amplitudes in some cells, but this fraction of cells (bars at x<0 in Figure S5) substantially decreased and disappeared as the noise amplitude increased.
2. Noise of a given amplitude had a more pronounced effect to increase firing frequency for slow-firing cells compared to moderate-firing and fast-firing cells (Figure 3 D-F).
3. In the fast-firing SAN cell population, noise decreased AP-firing rhythmicity. This was accompanied by a significant increase in the CV of AP firing frequency at noise amplitudes >25 pA (Figure 3G). An important and counterintuitive result was that noise application to slow-firing SAN cells improved AP-firing rhythmicity (CV decreased; Figure 3I).

**Figure 3.**
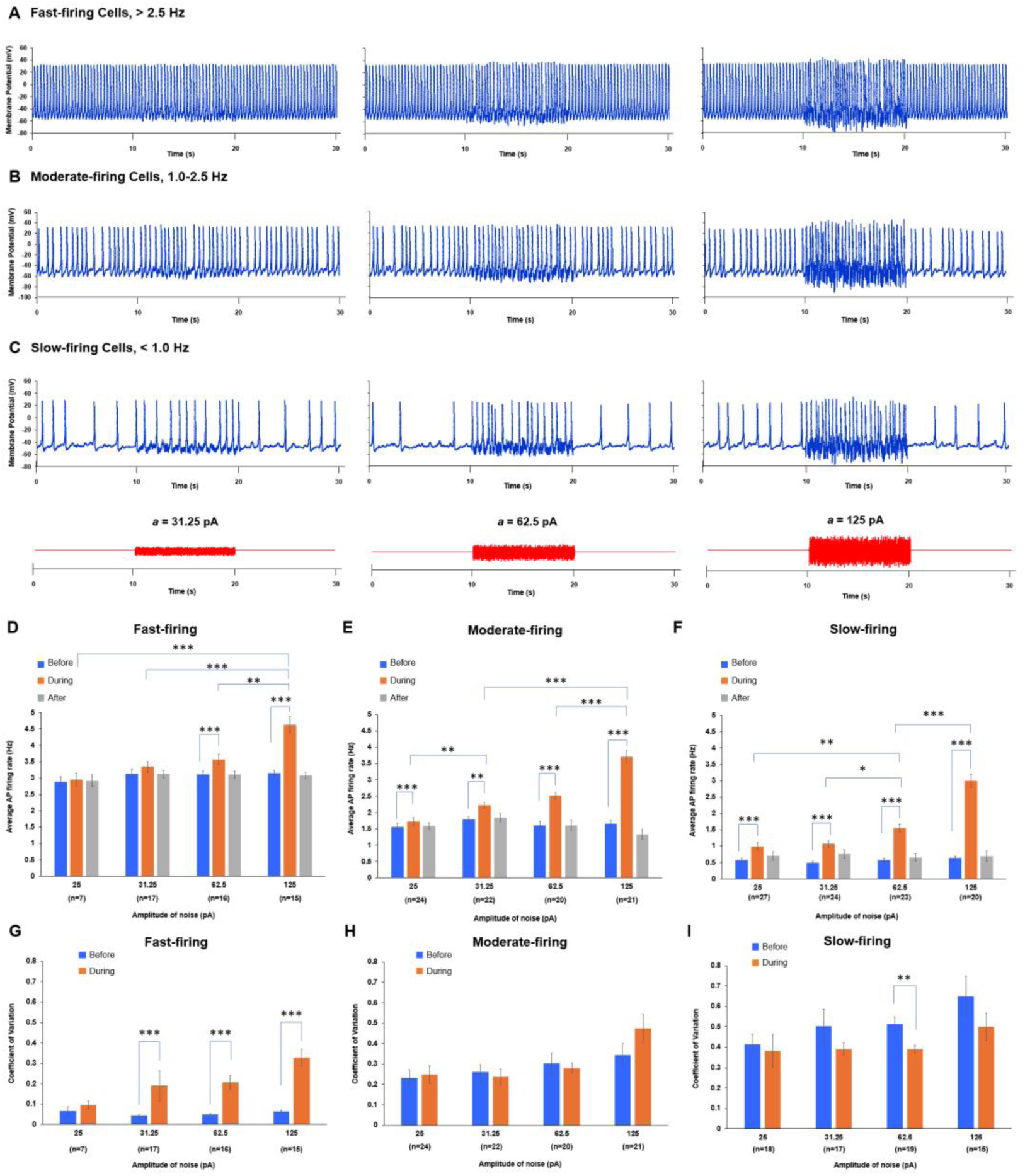
Stochastic resonance in SAN cell populations that spontaneously fired APs at different average rates. **A-C,** Examples of the effect of noise with different amplitudes (a) on AP firing in representative cells from each population. In all firing groups, noise increased AP firing rate, but it substantially perturbed the rhythmicity of fast-firing cells. D-F, Average AP firing rate was compared statistically before, during, and after noise application at each amplitude, and also among amplitudes (25, 31.25, 62.5, and 125 pA). In all cell populations, noise significantly increased AP firing rate (P < 0.05, repeated-measures ANOVA or Friedman test), except at 25 and 31.25 pA in the fast-firing group. There was no significant difference in AP firing rate between before noise application and after noise application was completed for any amplitude in any cell population. In the fast- and moderate-firing populations (**D and E**), 125 pA noise significantly increased the average AP firing rate compared with 25, 31.25, or 62.5 pA noise (p < 0.05, one-way ANOVA or Kruskal-Wallis test). In the slow-firing population (**F**), AP firing rate increased stepwise as noise amplitude increased. G-I, Similar statistical analyses were performed for the coefficient of variation (CV = SD/mean) of noise-induced AP firing. In the fast-firing SAN cell group (G), noise significantly decreased AP-firing rhythmicity (p < 0.05, paired t-test or Wilcoxon signed-rank test). In the moderate-firing group (H), there was no significant change in CV at any noise amplitude except 25 pA. In the slow-firing group (I), noise significantly decreased CV at 25 and 62.5 pA. *p < 0.05, **p < 0.01, ***p < 0.001.

### Stochastic resonance in dormant SAN cells

Subthreshold signaling during the first 10 seconds of recording (red boxed insets, Figure 4A) increased markedly when noise was applied from 10 to 20 seconds, eventually surpassing the AP threshold and culminating in AP firing. Figure 4B shows the distribution of AP firing rates across all dormant cells tested at each noise amplitude. Noise induced AP firing in every cell tested at 31.25, 62.5, and 125 pA. Even the smallest tested amplitude (25 pA) induced AP firing in most cells (21 of 24). The mean frequency of noise-induced APs increased significantly as noise amplitude increased (Figure 4C), consistent with the sine-wave results: a broader resonance spectrum at larger amplitudes (Figure 2L) allows processing of higher-frequency signals via stochastic resonance. Importantly, although firing frequency increased at higher amplitudes, rhythmicity decreased, and extremely strong noise abolished this benefit, with membrane potential (V_m_) showing high-amplitude fluctuations instead of APs (Figure S6). This behavior is consistent with stochastic resonance, which characteristically shows peak benefit at an optimal noise intensity.

**Figure 4.**
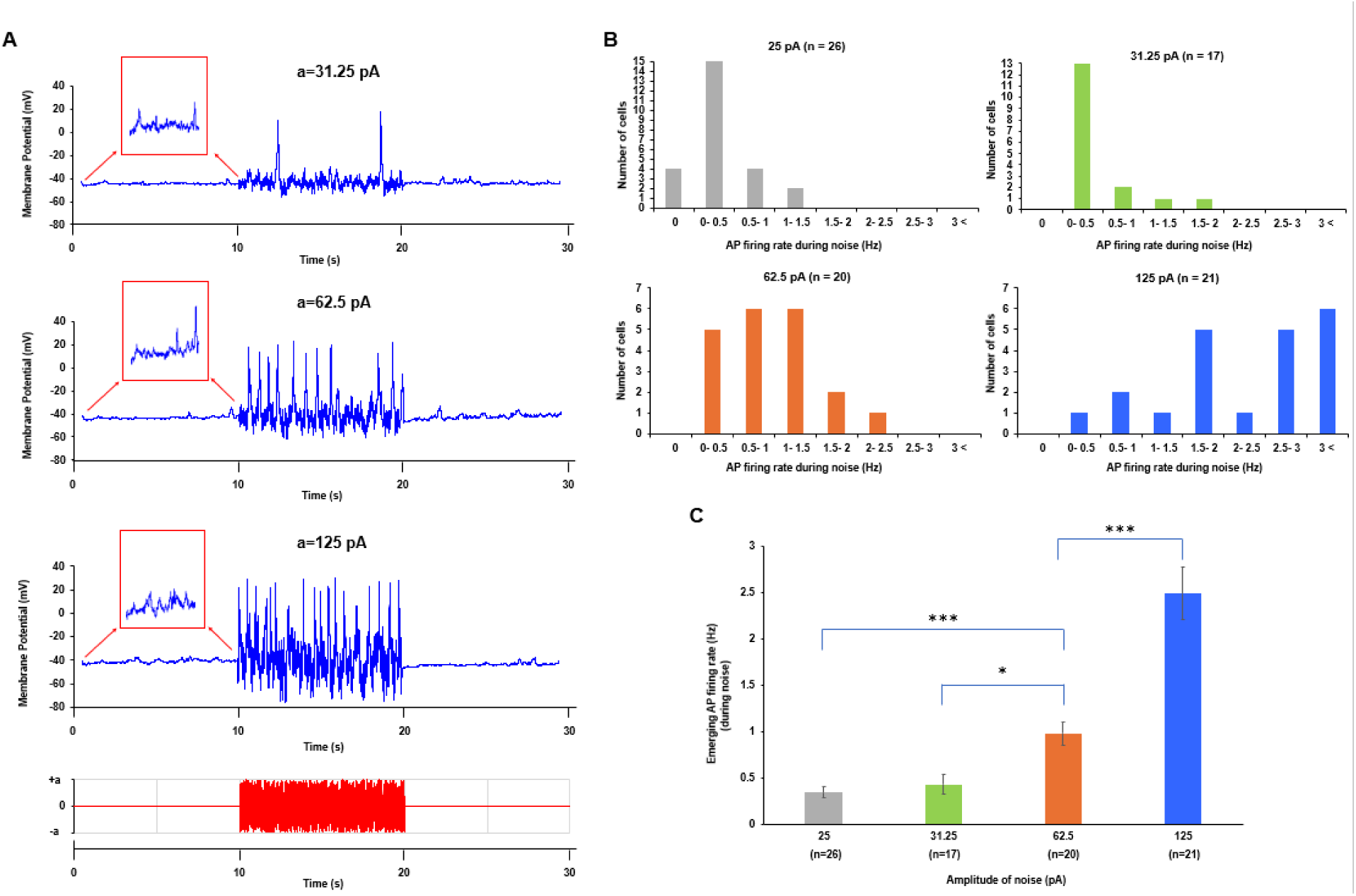
Stochastic resonance in dormant SAN cells. **A,** Representative examples of the effect of noise of different amplitudes on the AP firing rate. Before noise application dormant cells had subthreshold membrane potential oscillations (zoom-in, red squares) which were substantially increased in the presence of noise, reaching AP threshold and triggering AP firing, with the rate increasing as the noise amplitude increased. Moreover, the amplitude of noise-triggered APs substantially varied in each AP cycle, especially at larger amplitudes of noise. **B,** Distributions of the stochastic resonance effect for each noise amplitude tested. The noise awakened all dormant cells at noise amplitudes exceeding 25 pA. C, Average rate of AP firing (y-axis) caused by stochastic resonance increased as the noise amplitude increased (Kruskal-Wallis test). * p < 0.05, ** p < 0.01, *** p < 0.001.

### Resonance spectrum and stochastic resonance during cholinergic receptor stimulation

To decrease AP firing rate and ultimately induce dormancy in some AP-firing cells, carbachol (Millipore-Sigma 212385-M, a synthetic analog of acetylcholine) was applied consecutively at 100, 300, and 1000 nM. In cells that continued to fire in carbachol, the resonance spectrum shifted toward lower frequencies, as shown by responses to sine-wave stimulation. Figure 5A shows an example: in the basal state the cell fired spontaneously at about 4 Hz, whereas under carbachol it fired at about 0.33 Hz (3 APs in 10 s). Under carbachol the cell no longer fired APs at 5 or 6 Hz but responded rhythmically at lower rates such as 1 and 2 Hz (intervalogram in Figure 5B). These lower rates were not achieved in the basal state. As larger sine-wave amplitudes were applied in the presence of carbachol, the cell achieved higher AP firing rates (Figure 5C).

**Figure 5.**
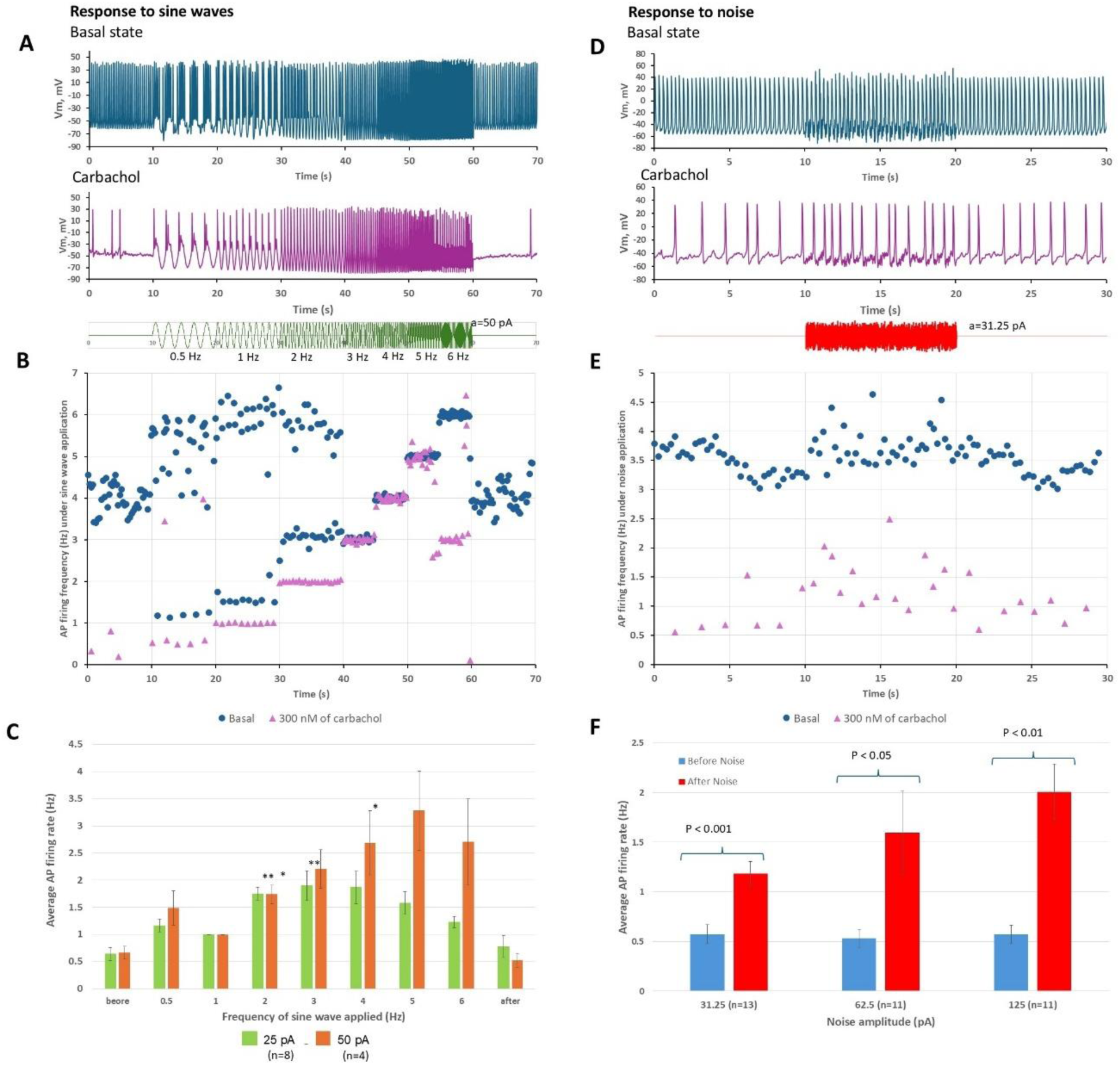
Resonance spectrum (functional range) and stochastic resonance in slow-firing cells during cholinergic receptor stimulation with carbachol. **A**, Original V_m_ recordings from the same cell in the basal state and in carbachol during the sine-wave protocol. The resonance spectrum shifted toward lower rates in the presence of carbachol. **B**, Intervalograms of the recordings in panel A. **C**, Statistical analysis of the effect of 25- and 50-pA sine waves on average AP firing rate (*p < 0.05, **p < 0.01, paired t-test for during-wave versus before-wave). **D**, Original V_m_ recordings from the same cell in the basal state and in carbachol during the noise protocol. Stochastic resonance increased AP firing rate in a representative slow-firing cell in the presence of carbachol. **E**, Corresponding intervalograms of the recordings in panel **D**. **F**, Statistical analysis of the effect of different noise amplitudes on averag**e** AP firing rate in cells under carbachol. The rate increased significantly at all amplitudes tested (P values are shown above the bars; paired t-test for during-noise versus before-noise).

Application of noise to firing cells under carbachol significantly increased their average AP firing rates (example in Figure 5D,E; population data in Figure 5F). As expected, an increase in firing rate is generally associated with lower AP rate variability, i.e. lower CV^34,35^. However, in our experiments a significant decrease in variability occurred only at a moderate (optimal) noise level. Thus, the benefit of rate improvement at stronger noise comes at the cost of greater rhythm disturbance, as large-amplitude noise introduces firing irregularity (Figure S7). This result is consistent with a hallmark of stochastic resonance: the presence of a maximum benefit at an optimal noise level.

Both sine-wave and noise protocols awakened dormant cells to generate APs (Figure 6). While a cell under carbachol began generating APs during sine-wave application, its maximum resonance frequency was reduced; for example, in panel A the maximum resonance frequency shifted from 2 Hz in the basal state to 1 Hz in carbachol (see red boxes in panel A and the intervalogram in panel B). A representative example of the noise effect is shown in panel D.

**Figure 6.**
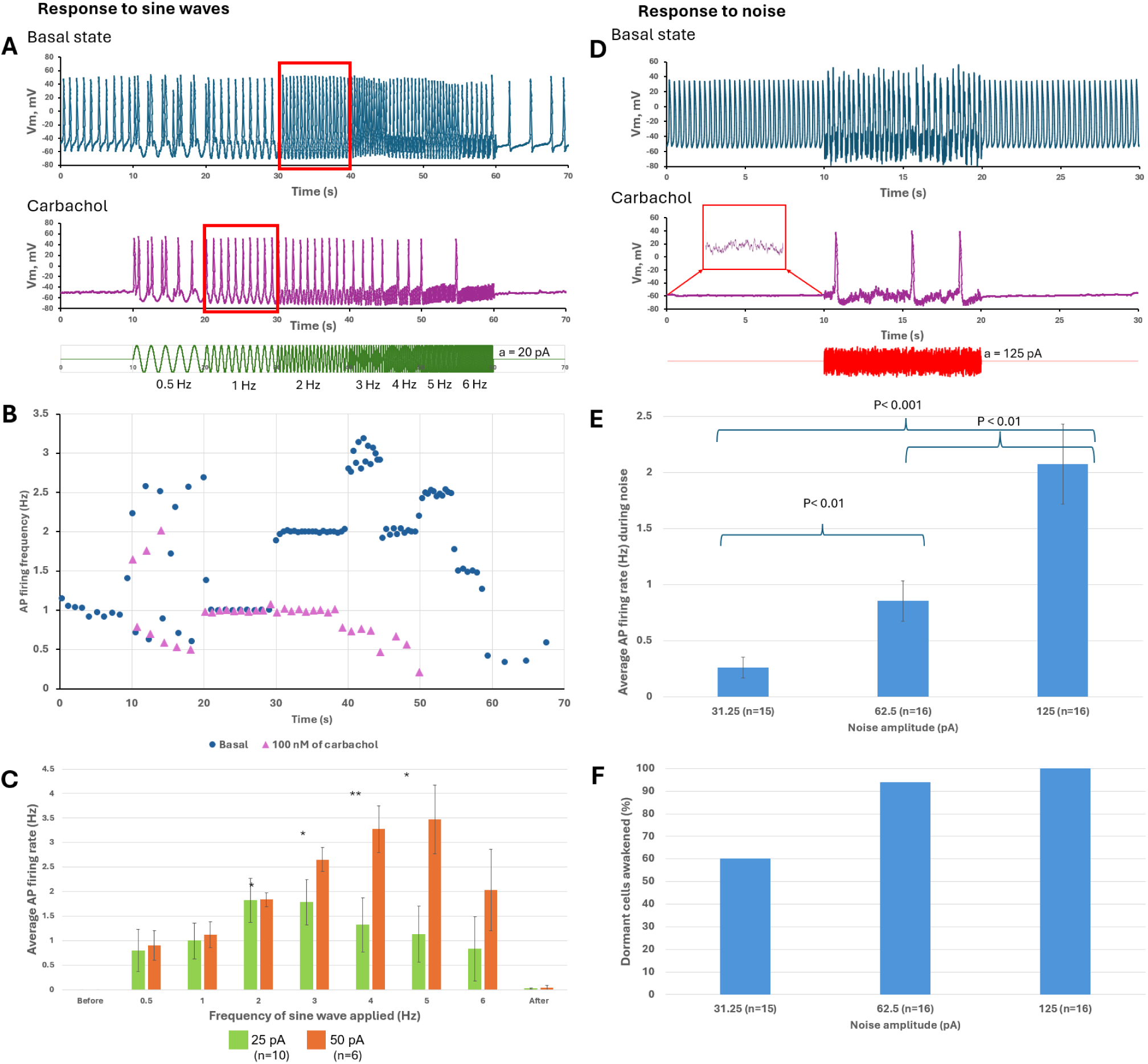
Resonance spectrum and stochastic resonance in dormant cells during cholinergic receptor stimulation with carbachol. **A**, Effects of a 20-pA sine-wave protocol in a representative cell in the basal state and after it became dormant in the presence of carbachol. The maximum resonance frequency shifted (red boxes) toward lower rates under carbachol (1 Hz) versus the basal state (2 Hz). **B**, intervalograms of recordings in **A**. **C**, Statistical analysis of the effect of sine waves of various frequencies and amplitudes (25 and 50 pA) on average AP firing rate (*p < 0.05, **p < 0.01, paired t-test for before-wave versus during-wave). **D**, Example of the noise effect on a dormant cell in the basal state and in the same cell during cholinergic receptor stimulation. The inset shows noisy intrinsic cell signals present under carbachol before noise was applied. **E**, Statistics for average AP firing frequency during noise at different amplitudes in the dormant-cell population. Statistical significance was determined by unequal-variance t-tests between pairs of different noise amplitudes (P values are indicated above each bar). **F**, Percentage of dormant cells awakened by a given noise amplitude.

Importantly, all dormant cells displayed noisy subthreshold V_m_ oscillation_s_ (red boxed inset in panel D), and these signals were amplified by noise to reach AP threshold via stochastic resonance. On average, larger sine-wave and noise amplitudes produced stronger effects, reflected in higher AP firing rates (panels C and E, respectively). Larger noise amplitudes also awakened a higher percentage of dormant cells (panel F). Thus, although cholinergic receptor stimulation shifts cell populations toward dormancy and slow firing, dormant cells can still be awakened to fire APs, and slow-firing cells can be induced to fire at higher rates, regardless of whether they occur naturally in the basal state or are induced by carbachol.

### Stochastic resonance is amplified by the coupled-clock system

We used three state-of-the-art numerical models, representing pacemaker function at three different scales (subcellular, cellular, and tissue), to support our experimental findings and gain insight into specific mechanisms of stochastic resonance in SAN cells. At the subcellular level, we employed our recent agent-based SAN cell model^36^ (parameters in the Supplemental Material), in which individual Ca release units (CRUs) form an interactive functional network via Ca-induced Ca release that is coupled to a full set of membrane currents. Consistent with our experimental results (Figure 4A), white-noise currents awakened a dormant cell model to begin firing APs, with firing frequency increasing at stronger noise amplitudes (Figure 7A).

**Figure 7.**
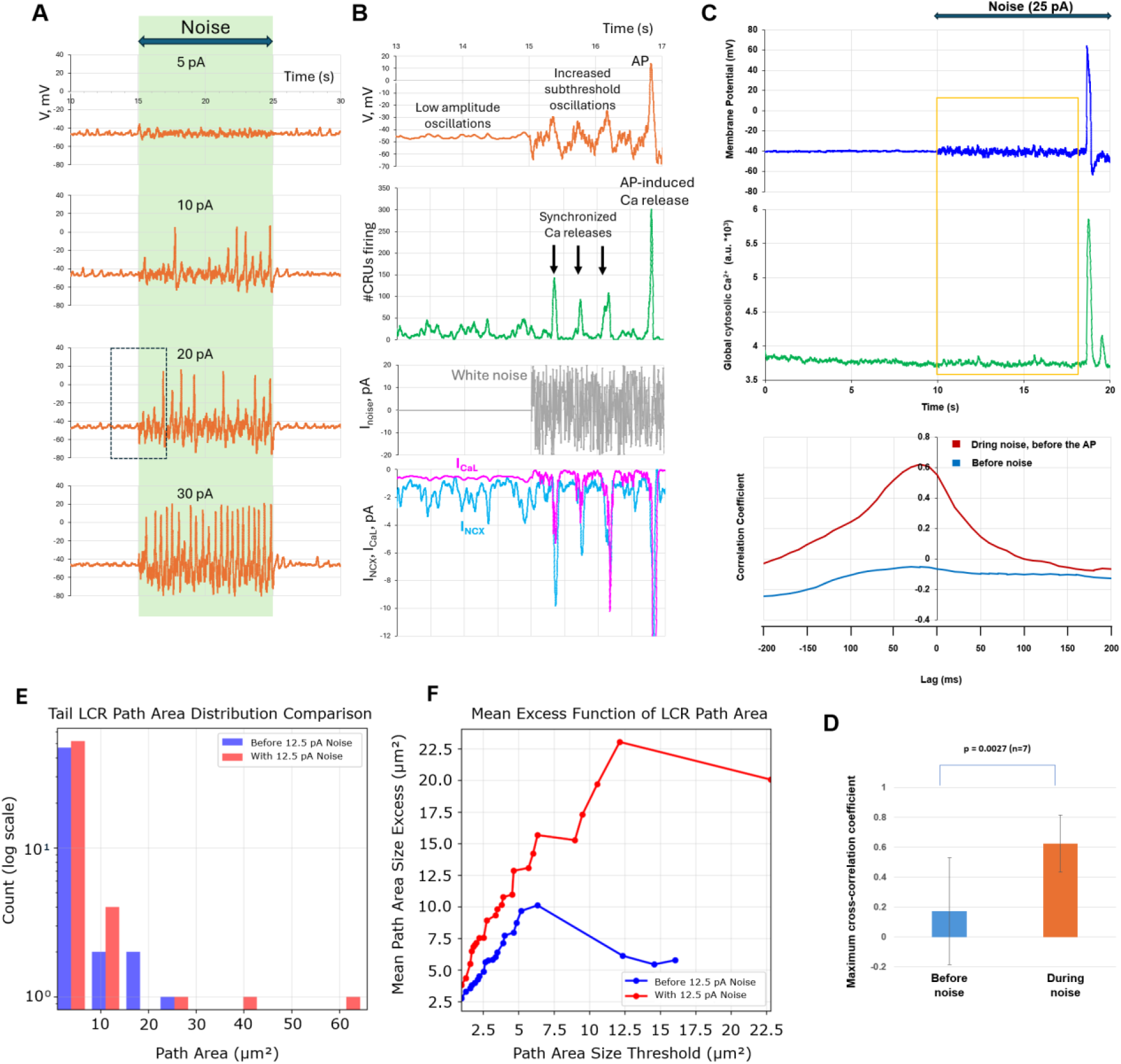
Stochastic resonance is amplified by the coupled-clock system in silico and in real cells. **A**, Responses of a dormant-cell model to noise of different amplitudes. **B**, Detailed comparison of cell signaling before and during 20 pA noise application before the first AP occurred (outlined by the box in panel A; see also Video S7). Synchronized CRU firing (green, downward black arrows), I_NCX_ (blue), I_CaL_ (magenta), and membrane-potential (V_m_) oscillations (orange) of increased amplitude, culminating in AP generation. **C**, Representative example of stochastic resonance in a representative dormant cell (see more examples in Figure S8) obtained in experimental simultaneous recordings of V_m_ and Ca signals (top panel), with the corresponding cross-correlation function for subthreshold V_m_ and Ca signaling before and during noise application (before the first AP fired). The time window chosen for cross-correlation analysis is shown by the orange square. **D**, Cross-correlation of subthreshold signals increased significantly during noise application, culminating in subsequent AP generation. **E**, Log**-**scale histogram revealing the architectural shift in LCR path areas between pre-noise (blue) and noise-present (red) states (cell #1 in Table S1). Noise exposure dramatically enriches the heavy-tail population of LCRs. **F**, To quantify heavy-tail behavior, large LCRs were further analyzed using mean-excess plots. Mean-excess analysis filters out LCRs above a given path-area threshold, calculates the average path area of the remaining larger LCRs, and compares this average with the original threshold. At higher cutoff levels, the average exceeds the threshold, confirming that large events become larger under noise conditions.

To gain insight into specific mechanisms of stochastic resonance in SAN cells, we compared subthreshold signaling before and during noise application in the dormant cell model. The amplitude of subthreshold V_m_ oscillations increased markedly during noise application because of coupled activation of CRUs, I_CaL_, and the Na/Ca exchanger current (I_NCX_), ultimately resulting in AP generation (Figure 7B). Ca releases became strongly synchronized in time and space and, as a result, increased in amplitude, with their peaks aligned with low-frequency inward-current components sensed and processed by the cell (resonance around 2 Hz; Video S7).

Experiments with simultaneous recording of V_m_ and Ca provided further evidence for subthreshold signaling amplification. The amplitude of the coupled subthreshold V_m_ and Ca fluctuations substantially increased in the presence of noise in the dormant cells (examples in Figures 7C, Figure S8, Video S8). To quantitatively demonstrate the increased coupling of Ca and V_m_ fluctuations, we examined their cross-correlations during noise (bottom sub-panels) and found that they significantly increased (Figure 7D). Importantly, to reveal genuine effects of subthreshold noise amplification, the cross-correlation was examined before the first AP was fired. Otherwise, the cross-correlation was always strong in AP firing cells independent of noise application, because each AP is strictly correlated with its attendant APCT (Figure S9).

To further explore how external noise amplifies cellular subthreshold signaling, we analyzed local Ca releases (LCRs) in dormant cells during simultaneous V_m_ and Ca recordings. Noise application in all analyzed cells (n=5) shifted LCR size distributions toward larger events quantified by a significant increase in L-kurtosis (mean 0.548 to 0.635; p<0.01; Table S1). LCR sizes were assessed as the entire Ca release propagation path area ^37^ reflecting CRU capability of recruiting neighboring CRU to fire via CICR mechanism. Under noise conditions, the LCRs exhibited a distinctly heavier tail (Figure 7E). Mean-excess analysis further confirmed that this shift reflected genuine enrichment of large-scale LCR events rather than uniform scaling (Figure 7F). This enrichment of large-scale LCRs reflects greater self-organization and synchronization of Ca release under noise.

### Stochastic resonance substantially expands the parametric space for AP firing in SAN cells

The “common-pool” Maltsev-Lakatta model^38^, which features faster computation, allowed us to test the effects of noise on coupled-clock function in a large number of derived cell models (n=23,668) representing different cell populations in the basal state and during cholinergic receptor stimulation. Experimental studies demonstrated that clock coupling determines the transition of a SAN cell from dormancy to rhythmic firing^14,16^. We therefore tested a wide variety of cell models with different basal-state firing activities generated by varying two key parameters of the coupled-clock system: I_CaL_ conductance (g_CaL_) and sarcoplasmic reticulum Ca pumping rate (P_up_). Variability in g_CaL_ reflects experimentally measured large-scale heterogeneity of I_CaL_ density^39–41^. Variability in P_up_ reflects a wide range of P_up_ inhibition levels such as arising from varying degree of phospholamban phosphorylation or pharmacological interventions. The diagram in Figure 8A shows the parametric space of firing cells in red shades, reflecting their average AP rates, and dormant cells in blue. Adding 25 pA noise awakened a major fraction of the dormant-cell population to fire APs, thereby extending AP firing (red area) toward the non-firing zone (blue area) of the parametric space in Figure 8B (see also Video S9). Representative examples of noise effects in the cell populations are shown in panels E-G: noise disturbed rhythmic firing in a fast-firing cell model, enhanced the rate and rhythm of a dysrhythmic slow-firing cell model, and awakened a dormant cell model to fire frequent APs, consistent with the experimental results (Figures 3A, 3C, and 4A). The stochastic resonance effect in awakening dormant cells was even more pronounced in the parametric space of SAN models during cholinergic receptor stimulation (Figure 8C,D; Video S9).

**Figure 8.**
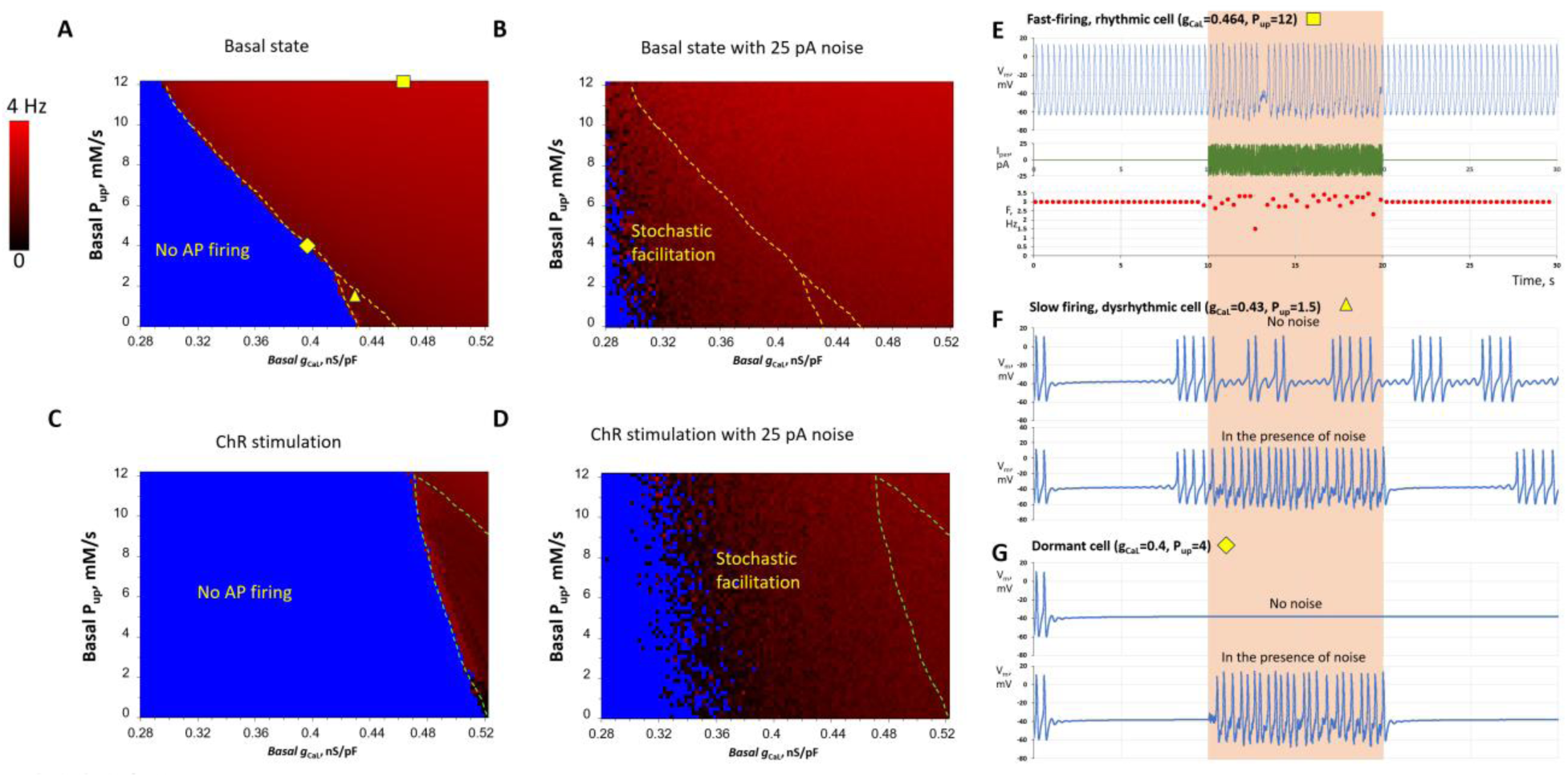
Insights from single-cell numerical modeling: stochastic resonance expands the parametric space of AP firing in the basal state and during cholinergic receptor stimulation. **A-D**, Each panel represents the result of numerical simulations of 5,917 (97 x 61 in xy) models with different coupled-clock parameters. I_CaL_ conductance (g_CaL_) was distributed evenly along the x-axis from 0.28 to 0.52 nS/pF in 0.0025-nS/pF increments. The sarcoplasmic reticulum Ca pumping rate (P_up_) was distributed evenly along the y-axis from 0 to 12 mM/s in 0.2-mM/s increments. Each pixel in the diagrams represents a single-cell model, with average AP firing frequency indicated by red shades. Blue areas indicate non-firing models. The blue area decreased substantially in the presence of noise. See Video S9 for V_m_ dynamics in all models. **E-G**, Representative examples of white-noise effects in different cell populations, with g_CaL_ and P_up_ values shown above the V_m_ traces in each panel. Noise disturbed the rhythmicity of AP firing in fast-firing cells but enhanced the function of slow-firing cells and awakened dormant cells to begin firing APs. The magenta double-headed arrow in panel A indicates the parameter range (g_CaL_=0.37 nS/pF, 0<P_up_<9.5 mM/s) used for the tissue-model simulations in Figure S12.

### Stochastic resonance prevents sinus arrest in a SAN tissue model

Parameters for the tissue model (Figure S12) were chosen so that most cells would lie in a non-firing zone, allowing the model to progress into sinus arrest after a few cycles over 6 s (Figure S12, Video S10). As previously demonstrated^42^, the transition to sinus arrest occurs via a phase transition in which dormant cells suppress the activity of all intrinsically firing cells. In the presence of noise, however, the SAN model continued to fire APs, clearly indicating that stochastic resonance prevents the phase transition of SAN tissue toward sinus arrest (Figure S12, Video S10).

## DISCUSSION

Recent high-resolution imaging studies portray a novel paradigm of SAN pacemaker function based on heterogeneous signaling within a brain-like cellular network^9,19^. The specific mechanisms driving pacemaking in such a complex network remain unknown. Noise plays a fundamental role in information processing and affects all aspects of nervous-system function^22,23^. One beneficial effect of noise in biological systems is stochastic resonance, which amplifies signaling in noisy environments and is used, for example, by neuronal networks for information processing^32^. Stochastic resonance models have also been conjectured for SAN function^17,18,29–31^. The present study provides the first direct experimental and theoretical evidence for the presence of stochastic resonance in SAN cells and its importance in pacemaker function.

### Specific mechanisms of stochastic resonance in SAN cells

While some SAN pacemaker cells beat spontaneously and are often called SAN myocytes, they share many properties with neuronal cells^43^. SAN cells express neuronal-type L-type Ca channels Ca_V_1.3^44^, neuronal-type Ca-activated adenylyl cyclases (AC1 and AC8)^45,46^, and the neuronal-type RyR3 Ca release channel^47^. They also share common properties with glutamatergic neurons^48^ and feature extended branches that support efficient long-range communication, as in neuronal networks. Pacemaker-cell interactions include pacemaker shift^49^, transitions to and from dormancy^10,12,14–16^, tonic entrainment^10^, and generation of complex signaling patterns, including rare firing and bursts of impulses^9,30^. Thus, SAN cells operate in a neuron-like manner: they detect, process, and respond to signals generated by other cells and, in turn, influence the operation of those cells.

The present study indicates that stochastic resonance is an important mechanism underlying these tasks. Depending on the signals processed via stochastic resonance, cells can shift their operating mode (i.e., firing modality^17^): a dormant cell can begin firing APs (Figure 4), a slow-firing cell can increase its firing rate (Figure 3), and cells can enter a burst-firing mode when low-frequency signals temporarily hyperpolarize the membrane (e.g., Figure 2A-C at 0.5 and 1 Hz). SAN cell AP firing can also depend on previous interactions, exhibiting a memory effect. For example, some dormant cells awakened by stochastic resonance continued firing APs even after noise removal (Figure 2H, group “After”; see also Figure S10). One possible mechanism is that cell activity becomes imprinted in altered states of sarcoplasmic reticulum Ca loading (short-term memory) or biochemical signaling (long-term memory), for example involving PKA, CaMKII, and adenylyl cyclases, which are critical for coupled-clock function in general^33^ and for restoration of AP firing in dormant cells in particular^14^. Thus, external signals can change the mode of SAN cell operation via stochastic resonance. The cell then responds by firing APs, sending signals to other cells in the collective SAN network for further information processing.

Signal processing and pacemaker-cell response depend on each cell’s resonance spectrum and its present operating status. The broadest resonance spectra were found in fast- and moderate-firing cells, whereas slow-firing and dormant cells exhibited narrower resonance spectra (orange bands in Figure 2A-D). Importantly, while resonance-spectrum width defines which frequency components can be utilized by a cell, the specific response to a complex perturbation (such as white noise in our experiments) depends on the cell’s operating point relative to AP threshold and on amplification of subthreshold V_m_-Ca oscillations. Therefore, dormant cells with a narrower low-frequency resonance spectrum can still undergo a qualitative transition from non-firing to firing (Figure 4), whereas among already firing cells the strongest rate facilitation was observed in slow-firing cells (Figure 3F).

Our numerical model simulations and simultaneous recordings of V_m_ and Ca revealed that stochastic resonance in SAN cells is not a simple electrical phenomenon: subthreshold signaling in SAN cells is strongly amplified by noise through coupled changes among local Ca releases, I_NCX_, V_m_, and I_CaL_ within the coupled-clock system (Figure 7, Figure S8, Videos S7 and S8). This powerful signal-amplification mechanism, involving positive-feedback interactions^50^, together with the broad resonance spectrum, makes stochastic resonance especially efficient in SAN cells for processing and responding to small subthreshold signals across broad frequency ranges.

In SAN tissue, such signals can be generated by other SAN cells operating at different frequencies and in different modes, by neuronal cells, and by other cell types (e.g., glial cells and S100B+/GFAP− interstitial cells^19^). All of these signals are, therefore, processed simultaneously in real time, analogous to analog computation in neurons^51^. Both stochastic resonance and the resonance spectrum of SAN cells are modulated by the autonomic system. For example, cholinergic receptor stimulation shifts them toward lower frequencies (Figures 5 and 6). As a result, the cells become unresponsive to higher-frequency signals but process lower-frequency signals more effectively.

### Stochastic resonance ensures fail-safe operation and prevents sinus arrest

In suppressing heart rate, the cholinergic receptor stimulation decreases the strength of diastolic pacemaker signals, with stronger stimulation culminating in SAN sinus arrest. In terms of evolution, the low-heart rate regime is extremely important for survival as it allows for energy conservation until most needed, such as during the fight-or-flight response. This regime must be balanced by powerful and robust signal processing mechanisms that prevent sinus arrest.

Stochastic resonance can be one of them, because bradycardic cholinergic receptor stimulation simultaneously increases overall biological noise in the SAN cells and tissue due to: 1) increased LCR stochasticity in SAN cells^52^; 2) an increased number of dormant cells^10^, that generate stochastic LCRs^12^; and 3) intensification of individual local releases of acetylcholine by nerve endings^25,26^. We show, in turn, that noise can stimulate dormant cells to fire APs (Figure 4), including those that become dormant during cholinergic receptor stimulation, and substantially enhance the rate and rhythm of cells firing infrequent, dysrhythmic APs (Figure 3). Furthermore, our numerical simulations indicate that these noise-mediated enhancements likely operate in a large fraction of SAN cells (Figure 8A-D and Video S9) and prevent sinus arrest in SAN tissue (Figure S12 and Video S10). In contrast, deterministic, limit-cycle models of SAN cells (lacking stochastic components) do not provide such robust function^53^.

Organisms harness stochasticity to generate a wide range of possible solutions to environmental challenges and to make optimal choices for future action^54^. In this sense, our study suggests that SAN cells and the SAN cellular network can employ stochastic resonance together with criticality-related behavior^2,55^ to expand the space of possible solutions, and thus the repertoire of complex responses, that help navigate future firing pattern and avoid AP cessation at low AP firing rates.

With respect to the fast-firing cell population, the effect of noise in these cells does not neatly fit the classic definition of stochastic resonance. These cells generate their own frequent rhythmic signals during each diastolic depolarization and reach AP threshold without further external amplification or prompting, whereas noise application leads to their dysrhythmic firing (Figure 3A,G). We also observed decreases in firing rate in some cells exposed to small and moderate noise amplitudes (Figure S5, bars at x<0; example in Figure S11), consistent with inverse stochastic resonance, in which noise suppresses oscillation frequency^56^. Such noise-induced suppression adds further complexity to the ways in which noise can regulate cell AP firing and, ultimately, heart rate. Thus, while noise is a universal modulator of SAN cell behavior, its role as a beneficial stochastic-resonance mechanism is most prominent at lower firing rates.

A recent conceptual model of stochastic resonance in the SAN^17,30^ has proposed that “noisy” cells with sporadic APs and subthreshold voltage fluctuations in the inferior SAN stabilize rhythm and prevent pauses, that is, prevent burst firing in the superior SAN. Our single-cell experiments partly support this hypothesis: if a single cell generates irregular bursts of AP firing, application of an optimal external noise can indeed eliminate the pauses (Figure 3B, middle panel). Furthermore, noise can promptly restore normal, rhythmic pacemaker function in populations of slow-firing cells (Figure 3C and 3F) and dormant cells (Figure 4). Our numerical model simulations demonstrate that stochastic resonance can improve pacemaker function at multiple scales across a wide range of key model parameters (Figures 7 and 8).

Previous studies in SAN tissue have shown that the central portion of the node actually beats slower than portions closer to the cristae terminalis when these pieces of the node are separated from each other^57,58^. Additionally, SAN pieces extracted near the left atrium were nonfunctioning until adrenaline or acetylcholine were applied. Taken together, these reports and our findings suggest that distinct clusters within the sinus node, characterized by varying pacemaking activity, may possess different sensitivities to stochastic resonance when integrated into a functional tissue network. For example, the SAN central pacemaking site may rely on stochastic resonance to maintain a faster beating rate. When isolated from the surrounding tissue, its rate may slow because it no longer receives noise input from neighboring cells. This reasoning is consistent with a recent experimental report that the cell cluster in the central SAN initiating the SAN impulse exhibits the most stochastic behavior^21^.

### New approaches to study the new pacemaker paradigm

In the classical paradigm of concentric excitation propagation within SAN tissue, emphasis was placed on parameters governing cell automaticity (I_f_), excitability (I_CaL_ and I_Na)_, and tissue conduction (connexins). In the emerging paradigm of information processing within the complex SAN cellular network^9,19^, additional parameters become important, including cell clusters forming brain-like small-world modular networks with efficient signal processing^11,59^; cell resonance spectra (Figure 2), which are critical for stochastic resonance efficiency; diversely firing cell populations^12,14,15^ (we studied four here); diversity of cell sensitivity to autonomic modulation^60^; phase-like transitions in which dormant cells prevail and the SAN abruptly falls into sinus arrest^42^; and locally released mediators that orchestrate cell activity and tune network operation. An extreme example of the importance of heterogeneity and clustering was recently demonstrated by two dormant cell populations that fired no APs in isolation but generated rhythmic APs when connected within a functional network^31^.

### Study limitations, future studies, and importance for aging and translational research

Although our experiments were performed in rabbit SAN cells, these cells share fundamental pacemaker mechanisms with human SAN cells ^16^, including coupled-clock operation, similar ion-channel expression, and heterogeneous cell populations featuring dormant cells. Critically, biological noise and background Ca signals analogous to those characterized here in rabbit SAN cells have been directly observed in human SAN tissue (Video 5 in Bychkov et al.^9^), supporting the translational relevance of our findings. The main species difference relevant to stochastic resonance is the mean heart rate, which would be expected to shift the resonance spectrum toward lower frequencies in humans. Further studies in human SAN cells will clarify how specific differences in ion channel densities, cell coupling, and autonomic innervation affect stochastic resonance characteristics.

Future studies in cells isolated from specific anatomical regions, as in^30^, and in new models such as brain-like small-world SAN models^59^, will examine how stochastic resonance operates within and among pacemaker cells, functional modules^9,11,30,31,59,61^, and other cell types in the SAN. Because stochastic resonance reflects a general ability of ion-channel systems to detect and amplify small signals in noisy environments^62^, and because threshold-dependent firing in noisy heterogeneous tissues is not unique to the SAN, future studies should also examine its role in other cardiac tissues, such as atrial and ventricular muscle, latent pacemakers in the atria, the AV node, and Purkinje fibers that generate escape rhythms when SAN function fails.

These questions are especially relevant to aging. Disorder within biological systems tends to increase with aging across scales, suggesting that heterogeneity and noise in SAN also increase with age. Aging-associated fibrosis, tissue fragmentation, and cell decoupling are likely to reduce transmission of noise between cells, but they can also increase the amplitude of local noise within individual cells by confining current fluctuations that would otherwise spread electrotonically to neighboring cells. Intermediate coupling supports SAN tissue beating through interactions between AP-firing and dormant cells^42,63^, whereas noise can be beneficial under partial decoupling, for example when ion-channel opening-closing noise increases SAN AP firing in Cx30-deficient mice^64^. Older SAN cells appear less able to generate and process higher-frequency signals^65^, consistent with the shift of the resonance spectrum and stochastic resonance toward lower frequencies that we observed during cholinergic receptor stimulation (Figures 5 and 6). Thus, stochastic resonance likely becomes more important in supporting function of the aged SAN at low rates, partially compensating for the reduced intrinsic heart rate and helping prevent sinus arrest.

Sick sinus syndrome associated with bradyarrhythmia and sinus arrest remains a major healthcare problem, especially as the elderly population grows^3^. Furthermore, sinus node dysfunction affects up to one in five patients with atrial fibrillation^66^ and is also associated with severe bradyarrhythmia in heart failure^67^, which can lead to sudden cardiac death. Our findings may help develop therapeutic strategies that, rather than relying solely on direct pacing, restore or substitute for complex cellular signals that are lost or degraded with aging, thereby treating bradyarrhythmia and preventing sinus arrest.

Another potential avenue is the development of biological pacemakers. Although this idea was proposed more than two decades ago^68,69^, progress has been limited, in part, by incomplete understanding of the fundamental principles governing SAN cellular-network operation^2^. The emerging paradigm of cardiac pacemaker function^2,9,10,17–19,29–31^, with stochastic resonance as a key component, suggests that bioengineering strategies should be revisited on a new conceptual basis to create biological pacemakers that more faithfully mimic the native heart.

## Acknowledgments

The authors acknowledge the assistance of Bruce D. Ziman for skillful isolation of sinoatrial node cells.

## Supplemental Methods

### Rabbit SAN cell isolation

SAN cells were isolated from male rabbits as previously described^70^, in accordance with NIH guidelines for the care and use of animals (protocol #457-LCS-2024). New Zealand White rabbits (Charles River Laboratories, USA) weighing 2.8–3.2 kg were anesthetized with sodium pentobarbital (50–90 mg/kg). The heart was removed quickly and placed in a solution containing (in mM): 130 NaCl, 24 NaHCO_3_, 1.2 NaH_2_PO_4_, 1.0 MgCl_2_, 1.8 CaCl_2_, 4.0 KCl, and 5.6 glucose, equilibrated w_i_th 95% O_2_ / 5% CO2 (pH 7.4 at 35°C). The SAN region was cut into small strips (∼1.0 mm wide) perpendicular to the crista terminalis and excised. The final SAN preparation, consisting of SAN strips attached to a small portion of the crista terminalis, was washed twice in nominally Ca-free solution containing (in mM): 140 NaCl, 5.4 KCl, 0.5 MgCl_2_, 0.33 NaH_2_PO_4_, 5 HEPES, and 5.5 glucose (pH 6.9) and incubated on a shaker at 35°C for 30 min in the same solution supplemented with elastase type IV (0.6 mg/mL; Sigma Chemical Co.), collagenase type 2 (0.8 mg/mL; Worthington, NJ, USA), protease XIV (0.12 mg/mL; Sigma Chemical Co.), and 0.1% bovine serum albumin (Sigma Chemical Co.). The SAN preparation was then placed in modified Kraftbruhe (KB) solution containing (in mM): 70 potassium glutamate, 30 KCl, 10 KH_2_PO_4_, 1 MgCl_2_, 20 taurine, 10 glucose, 0.3 EGTA, and 10 HEPES (titrated to pH 7.4 with KOH), and kept at 4°C for 1 h in KB solution containing 50 mg/mL polyvinylpyrrolidone (PVP; Sigma Chemical Co.). Finally, cells were dispersed from the SAN preparation by gentle pipetting in KB solution and stored at 4°C.

### Electrophysiology and Ca signal imaging

Perforated patch-clamp recording was used in current-clamp mode to measure V_m_ and apply external membrane currents of different waveforms. Membrane patch perforation was achieved using β-escin (50 μM) added to the patch-pipette solution, as described elsewhere ^71^. The patch-pipette solution contained (in mM): 120 K-gluconate, 5 NaCl, 5 Mg-ATP, 5 HEPES, 20 KCl, and 3 Na_2_ATP (pH adjusted to 7.2 with KOH). Axopatch 200B, DIGIDATA 1440, and pCLAMP software (Molecular Devices, USA) were used for data acquisition with a sampling interval of 0.1 ms.

Ca imaging was performed as previously described ^14^. Cells were loaded with 5 μM Fluo-4AM for 20 min at room temperature before measurement. Ca signals were imaged with a 2D sCMOS camera (PCO edge 4.2) with a 13.2-mm square sensor and 2048 x 2048 pixel resolution. Images were acquired at 100 frames/s, which required use of only part of the sensor (1280 x 1280 pixels). The recording camera was mounted on Zeiss Axiovert 100 inverted microscopes (Carl Zeiss, Inc., Germany) with a 40x oil-immersion lens and a CoolLED pE-300-W fluorescence excitation light source (CoolLED Ltd., Andover, UK). Fluo-4 fluorescence was excited at 470/40 nm and emission was collected at 525/50 nm using Zeiss filter set 38 HE.

Measurements of V_m_ separately or simultaneously with Ca were performed at 36°±0.1°C (500 μl chamber volume). Temperature was controlled by an Analog TC2BIP 2/3Ch bipolar temperature controller from CellMicroControls (Norfolk, VA, USA). This heated both the glass bottom of the perfusion chamber and the solution entering the chamber (via a pre-heater). The physiological (bathing) solution contained in mM: NaCl 140; KCl 5.4; MgCl_2_ 2; HEPES 5; CaCl_2_ 1.8; pH 7.3 (adjusted with NaOH). In order to perform simultaneous recording of Ca signals and membrane potential, we programmed the PCO camera software to generate a TTL signal when imaging started. This signal triggered V_m_ recording via pCLAMP software. Video-recordings of intracellular Ca signals were analyzed by ImageJ program.

### Cell populations tested in the study

We tested four cell populations based on intrinsic activity: (i) fast-firing cells (>2.5 Hz, i.e., classical pacemakers); (ii) moderate-firing cells (1 to 2.5 Hz); (iii) slow-firing cells (<1 Hz); and (iv) dormant cells firing no APs. Notably, dormant cells had morphology similar to that of classical pacemaker cells, being mainly spindle-shaped (Figure S2). Dormant cells were either (i) naturally dormant, that is, generating no APs at zero current clamp; (ii) rendered dormant by application of carbachol, a synthetic analog of ACh; or (iii) rendered dormant by passage of a hyperpolarizing current simulating electrotonic interactions with neighboring cells, such as the influence of other dormant cells ^10^ or cells near the atrial tissue surrounding the SAN.

### Mouse heart isolation

The mouse heart was dissected as previously described ^9^. Experimental protocols were approved by the Animal Care and Use Committee of the National Institutes of Health (protocol #034-LCS-2019). The heart was removed quickly and placed in standard Tyrode solution containing (in mM): 130 NaCl, 24 NaHCO_3_, 1.2 NaH_2_PO_4_, 1.0 MgCl_2_, 1.8 CaCl_2_, 4.0 KCl, and 5.6 glucose, equilibrated with 95% O2 / 5% CO2 (pH 7.4 at 35.5°C).

### Preparation of mouse SAN tissue and Ca signal imaging by high-speed camera

Preparation of mouse SAN tissue has been described in detail in ^9^. In short, the whole heart was pinned to a silicon platform under a surgical microscope in order to excise the right and left atria. A 10-ml tissue bath was perfused with standard solution at a rate of 10 ml/min. After removal of the ventricles, the right atrium was opened to expose the crista terminalis, the inter-caval area, and the inter-atrial septum. The preparation was not trimmed, leaving SAN region together with surrounding atria and superior and inferior vena cava (SVC and IVC) intact. The SAN preparation was pinned to the silicon bottom of the experimental chamber by small stainless-steel pins with the endocardial side exposed. Care was taken to provide the minimal amount of stretch required to flatten the SAN tissue. After mounting, the preparation was superfused with solution maintained at a temperature of 36±0.3°C.

We used the imaging system described previously ^9^ to assess intracellular Ca dynamics within individual cells across the intact mouse SAN. In brief, we used a stationary fixed-stage upright microscope (AxioExaminer D1 equipped with a zoom tube [0.5–4x], Carl Zeiss Microscopy LLC) and a PCO edge 4.2 camera featuring a scientific complementary metal-oxide semiconductor (sCMOS) sensor with high spatial and temporal resolution. The experimental chamber containing the SAN preparation was placed on a platform (Sutter Instruments) mounted on a pressurized air table (Newport).

The SAN preparation was incubated with a membrane-permeable Ca indicator Fluo-4AM (10µM) for 1.5 hours. The excitation light was generated by CoolLED pE-300ultra. The excitation light was reflected to the SAN preparation by a dichroic mirror with a central wavelength of 498 nm, and the emitted fluorescence signal was collected through a 530±20 nm filter (Semrock, USA). The fluorescence image of the SAN preparation was projected by air or water lenses onto the sCMOS camera sensor. To prevent interference of tissue motion during recordings from SAN tissue, we decoupled electrical excitation and mechanical contraction in some preparations by inhibiting the formation of the Ca-sensitive regulatory complexes within sarcomeres using 10 µM cytochalasin B ^72^.

### Preparation of mouse SAN tissue and Ca signal imaging by confocal microscopy

The heart was transferred to oxygenated Tyrode solution (35 ± 0.5°C) containing (in mM) 140 NaCl, 5.4 KCl, 1.2 KH_2_PO_4_, 1.0 MgCl_2_, 1.8 CaCl_2_, 5.55 glucose, and 5 HEPES, with pH adjusted to 7.4 with NaOH. It was pinned onto a silicone dish, the right atrium was opened, and the SAN region, including the superior and inferior venae cavae (SVC and IVC), was dissected and pinned with the endocardial side facing upward on a silicone ring. Calbryte 520 AM (20 μM) was added to the SAN preparation in oxygenated Tyrode solution, and the preparation was left for ∼2 h at room temperature. After 2 h, the preparation was washed and transferred to the experimental chamber, where the silicone ring was turned upside down (SAN endocardial side facing downward) for data acquisition (35 ± 0.5°C). Ca signals were recorded in Tyrode solution using a 10x/0.3 objective with an LSM 510 META confocal microscope (Carl Zeiss) in frame mode with 488-nm excitation. The pinhole was ∼4.2 Airy units; the spatial scale was ∼0.21–0.48 μm; the stack size was ∼108–245 μm; scan zoom was ∼3.7–8.3; and laser power was ∼5%–15%.

### Statistics

Values are expressed as mean ± standard error. Normality was tested with the Shapiro-Wilk test. Normally distributed continuous variables were compared using the paired t-test. One-way analysis of variance (ANOVA) with Bonferroni correction, repeated-measures ANOVA when appropriate, and the Wilcoxon signed-rank, Kruskal-Wallis, or Friedman tests for skewed data were used as appropriate. Trends in average AP firing rate during sine-wave application were assessed with the Jonckheere-Terpstra trend test. A value of p < 0.05 was considered statistically significant.

### Methods for LCR signal detection in isolated SAN cells

LCR analysis was performed using a custom Python pipeline that implements and extends the computational framework for LCR detection described in our previous work^37,73^. To ensure valid paired comparisons of LCR characteristics, equal-length time windows were extracted before and during noise application.

When TIFF image stacks are first loaded, the signal intensity is globally normalized by min-max scaling. Then to delineate the cell cytoplasm from the background, a binary cell mask is generated by creating a maximum-intensity projection over time and thresholding the resulting image at its mean intensity. The largest resulting connected component is retained as the final cell mask. Global fluorescence decay and drift are removed by subtracting the mean intensity within the cell mask from each frame and any negative differences are set to zero to produce a detrended image stack.

To emphasize the rising phase of Ca signals, frame-to-frame differences of the detrended signal were computed, and negative differences were again set to zero. Unlike our previous single-threshold approach^37^, which applies a one-pass cutoff based on the standard deviation (σ) of fluorescence intensity within the cell region to baseline-subtracted fluorescence, we applied a novel two-stage hysteresis thresholding procedure to these temporal differences to yield more robust, noise-resilient detection of true Ca release events. First, an initial seed threshold at sd_seed x σ (sd_seed = 1.0) identified core release areas; next, those cores were grown by including connected pixels above a lower growth threshold at sd_grow x σ (sd_grow = 0.5). Finally, candidate regions underwent connected-component labeling, size filtering, and morphological operations (closing, opening, and hole filling) to produce the final binary LCR mask for each frame.

Finally, LCR events were reconstructed into full spatiotemporal trajectories by linking regions across consecutive frames on the basis of maximal pixel overlap, a process that resolves LCR collisions and logs births, branches, and deaths, as described previously^37^. From these complete trajectories, per-event metrics were extracted, including path area, lifetime, and initiation/termination times, for subsequent statistical analysis. Our Jupyter Notebook for the full LCR analysis is available on GitHub (https://github.com/alexmaltsev/SANC-LCR-Analysis).

### Algorithm for background signal detection in SAN tissue

To better observe and analyze LCRs and other intrinsic noise patterns in SAN tissue, we developed a signal-enhancement pipeline centered on Penalized Matrix Decomposition (PMD), implemented through the Python package Trefide ^74^. Originally designed to extract neuronal Ca and voltage signals from brain-imaging data, this algorithm is well suited to analysis of complex spatiotemporal Ca dynamics in cardiac tissue, including the stochastic signals relevant to pacemaker function.

The pipeline begins with a custom automated preprocessing sequence to delineate the active tissue area in the raw TIFF movie (Figure S1A). First, the video’s contrast is enhanced using histogram normalization, where the intensity range is set by saturating 0.35% of the brightest and dimmest pixels. Next, a Gaussian Mixture Model (GMM) is fit to the maximum intensity projection to automatically define a tight bounding box around the SAN tissue. The raw video is then cropped to this box, and any remaining pixels whose maximum intensity across all frames fall below the mean maximum intensity of the field of view are masked and set to zero.

Following preprocessing, the prepared imaging data undergoes signal enhancement through PMD as implemented in Trefide (Figure S1B). Within each spatial patch (40×40 pixels), the algorithm identifies and extracts distinct signal activity patterns through a constrained optimization process that iteratively finds spatiotemporal components maximizing data variance while enforcing spatial coherence and temporal smoothness, capturing up to 50 separate signal sources per patch. In SAN tissue, these spatiotemporal components correspond to synchronized Ca transients within the tissue (which are later removed in further processing), propagating Ca waves, or localized Ca release events.

The decomposition ensures biological plausibility by applying mathematical constraints: a spatial penalty (Total Variation) maintains connected regions rather than scattered pixels, while a temporal penalty (Trend Filtering) preserves smooth Ca dynamics with sharp upstrokes characteristic of Ca events. To prevent the extraction of spurious patterns from measurement noise, the algorithm automatically terminates when it fails to identify statistically significant signal patterns in three consecutive attempts.

After processing all patches independently, Trefide reconstructs a complete enhanced movie by combining the extracted Ca signals through weighted averaging in overlapping regions, where signals present in multiple patches are blended based on their proximity to patch centers. This results in dramatically improved visualization of Ca activity, revealing previously obscured Ca dynamics including the stochastic Ca release patterns essential for understanding how SAN cells harness biological noise to ensure rhythmic heartbeat generation. Our Jupyter Notebook for the full SAN analysis is published on GitHub. (https://github.com/alexmaltsev/SAN-Analysis)

Following signal enhancement, Principal Component Analysis (PCA) removes the dominant global Ca transient from underlying local signals (Figure S1C). This approach draws from random matrix theory applications to cardiac tissue, as demonstrated by Norris & Maltsev ^11^, who showed that the first principal component of SAN Ca imaging data captures the collective, action potential (AP)-driven Ca wave propagating across the tissue. Our implementation follows a similar algorithm: it reshapes the signal-improved 3D movie into a 2D matrix (time × pixels), performs Singular Value Decomposition (SVD) to compute the first principal component, and subtracts it from the post-Trefide processed data. The resulting residual movie preserves local Ca signaling patterns while removing the dominant global transient.

Finally, to create an objective final mask of LCRs and other intrinsic noise patterns in SAN tissue, we used statistical methods from Extreme Value Theory (EVT) to derive a nonarbitrary, data-driven threshold for separating events from background fluctuations (Figure S1D). After subtraction of the global PCA component, most pixel residuals collapsed around zero, leaving approximately the remaining <10% of variance ^11^ split between small background fluctuations and occasional large-amplitude Ca transients which, by virtue of their magnitude and limited spatial footprint, form the extreme right-hand tail of the residual-intensity distribution. Thus, an optimal gray threshold could be computed to determine where heavy-tail events begin.

To determine this threshold, the algorithm fit a Generalized Pareto Distribution (GPD) to data values exceeding 100 different thresholds spanning quantiles from the 50th to 99th percentile. For computational efficiency, fitting proceeded only when at least 50 pixels exceeded a given threshold, and the data were downsampled to a maximum of 10,000 points. The key insight lies in analyzing the stability of the GPD shape parameter (ξ) as a function of threshold: the optimal threshold occurs where this parameter interpolates to zero. Applying this EVT-derived threshold to clip the residual movie effectively isolates LCRs as the remaining signals for subsequent analysis (Figure 1 and Figure S1E; see also Videos S1 and S2).

### Computer code and specific model parameters used in the present study

A key part of our study was numerical model simulations that supported our experimental results and provided further insights into how specifically stochastic resonance contributes to pacemaker function at three scales: (i) the subcellular scale, using Ca-Release-Unit (CRU)-based models^36^; (ii) cellular scale, using common pool models^38^; and (iii) SAN tissue scale, using a 2D grid multi-cellular model^42^.

### Code availability

We employed three numerical models at different scales:

1. Single SAN cell model at the subcellular level was our CRU-based (agent-based) model ^36^. Its computer code (Delphi language) is freely available as supplementary data to our original publication^36^. Code for more recent model versions is also freely available on GitHub: https://github.com/victoramaltsev/CRU-based-SANC-model (with various CRU sizes ^75^) and https://github.com/victoramaltsev/RyR-network-SANC-model (with individual RyRs ^76^).
2. Single SAN cell model at the cellular level (common-pool model) was the Maltsev-Lakatta model ^38^. The computer code for the original model is freely available in CellML format (maltsev_2009_paper.cellml) at http://models.cellml.org/workspace/maltsev_2009 and can be executed using the Cellular Open Resource (COR) software developed at the University of Oxford by Garny et al.^77^ (for recent development of COR, see http://www.opencor.ws).
3. The SAN tissue model used here is an adapted version^42^ of the tissue model originally developed by Campana. The model code in CUDA C is freely available in Campana’s PhD thesis, Appendix A^78^: https://amslaurea.unibo.it/8596/1/campana_chiara_tesi.pdf

### Parameters of noise

The noise current I_noise_ was generated as a sequence of random numbers within a range of [-a, a] with an interval of 4 ms (Figure S3). Intrinsic membrane current I_m_ together with I_noise_ drives V_m_ change: dV_m_/dt = - (I_m_+I_noise_)/C_m_, where C_m_ is cell membrane capacitance in the tested models and in real cells.

### Parameters of CRU-based model (agent-based model)

The full set of model parameters and computer code are given in our original publication ^36^. The dormant-cell model tested in the present study had the following specific parameters: CRU positions were distributed uniformly at random, and g_CaL_ was 0.27 nS/pF (see Figure 7B in ^36^).

### Important note about cell membrane capacitance

SAN pacemaker cell sizes are extremely heterogeneous, with cell membrane capacitance varying from 10 to 100 pF ^39,41,79^. Hence, the same current produces a stronger effect in smaller cells, because the V_m_ change dV during time period dt is directly proportional to the current dI but inversely proportional to the membrane capacitance Cm: dV=- dI*dt/Cm. Our model simulates function of a smaller (but still realistic) SAN cell having a membrane electrical capacitance of 20 pF. We chose such cell in our numerical model simulations because it better reproduces function of so-called “central cell” located in the SAN center where the leading pacemaker site is located. The cells in the center of the SAN are characterized by smaller size vs. peripheral cells ^80,81^. A 20 pF cell model has been previously used by other scientists to simulate function of the central cell ^82^. Thus, for example, in our 20 pF cell model current 20 pA noise (density 1 pA/pF) would produce similar V_m_ change as 62.5 pA current in a 62.5 pF cell (same density of 1 pA/pF).

### Parameters of Maltsev-Lakatta models (common pool models)

The full set of model parameters is given in our original publication ^38^. To generate models representing different cell populations, we varied two key parameters of the coupled-clock system, g_CaL_ and P_up_. Specifically, the model sensitivity analysis (Figure 8A-D) was performed by varying g_CaL_ from 0.28 to 0.52 nS/pF in 0.0025-nS/pF increments and P_up_ from 0 to 12 mM/s in 0.2-mM/s increments. In panels E, F, and G of Figure 8, the fast-firing cell was modeled with g_CaL_ = 0.464 nS/pF and P_up_ = 12 mM/s (the standard Maltsev-Lakatta model), the slow-firing cell with g_CaL_ = 0.43 nS/pF and P_up_ = 1.5 mM/s, and the dormant cell with g_CaL_ = 0.4 nS/pF and P_up_ = 4 mM/s.

The effect of cholinergic receptor stimulation with 0.1 μM acetylcholine was modeled as previously described^59^. In brief, the fractional block (b_CaL_) of I_CaL_ during cholinergic receptor stimulation was adopted from the model of Zaza et al.^83^, as given in the legend to their Figure 2:

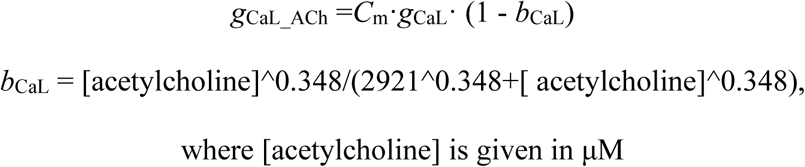

Thus, the fractional block in the presence of 0.1 μM acetylcholine (simulated in the present study) was relatively small: b_Ca*L*_(0.1) = 0.02728, that is, <3%. The shift s (in mV) of the I_f_ activation curve during cholinergic receptor stimulation was adopted from Zhang et al.^84^ as follows:

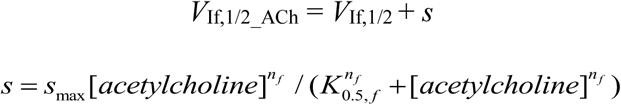

*s*_max_ = -7.2 mV: maximum acetylcholine-induced shift of *I*_f_ half activation voltage.

*n_f_* = 0.69 and *K_0.5,f_* = 12.6 nM: Michaelis-Menten parameters for acetylcholine modulation of *I*_f_. For 100 nM of acetylcholine: *s_max_* =-5.81 mV and *V*_If,1/2_ACh_ = -64 - 5.81 = - 69.81mV

The formulation of the acetylcholine-activated K current (I_KACh_) was adopted from Demir et al.^85^ (note that I_KACh_ = 0 when [acetylcholine] = 0), with g_KACh_ = 0.14241818 nS/pF.

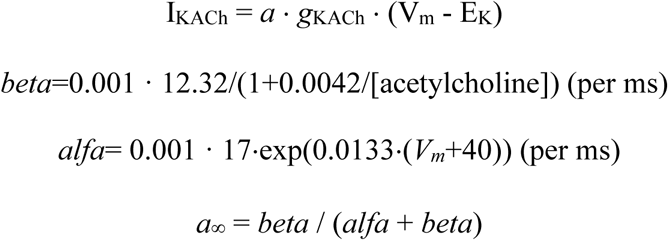

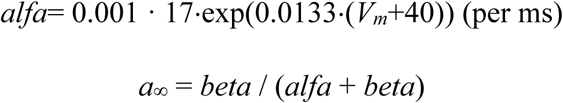

The present study did not investigate I_KACh_ kinetics or system transitions after acetylcholine application. Therefore, in our simulations I_KACh_ was assumed to have reached steady state, with a set to the steady-state value defined above. Thus,

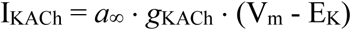

cholinergic receptor stimulation inhibits P_up_, with fractional block *b*_up_ formulated as follows:

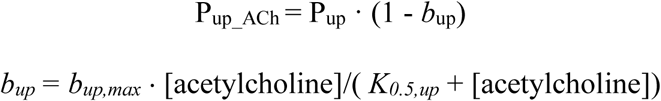

where *K_0.5,up_* = 90 nM is the [acetylcholine] for half-maximal inhibition and *b_up,max_* = 0.7. For 100 nM of acetylcholine: *b_up_ =*0.368421052632.

### Parameters of SAN tissue models (2D grid multi-cell model)

The full set of model parameters is given in our original publication ^42^. To generate the sinus-arrest model examined here, we distributed tissue-model parameters over the range g_CaL_ = 0.37 nS/pF and 0 < P_up_ < 9.5 mM/s (magenta double-headed arrow in Figure S12) so that most cells were dormant and the model progressed into sinus arrest after a few cycles over 6 s.

## Supplemental Figures S1-S12

**Figure S1.**
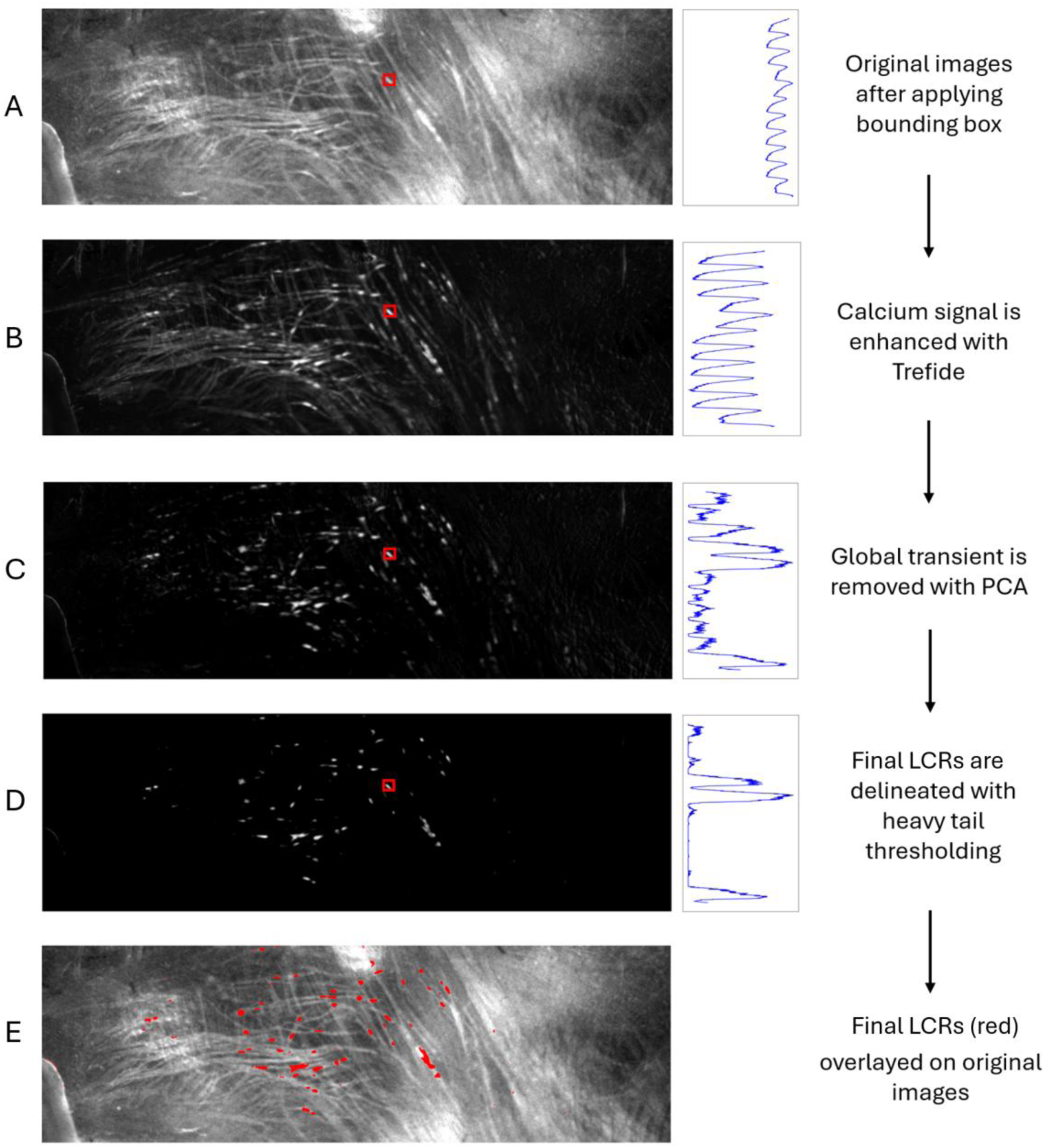
Step-by-step breakdown of the analysis used to identify LCR signals in SAN tissue. **A**, Preprocessed Ca-imaging data following automated tissue-boundary delineation and spatial cropping, showing raw fluorescence intensity after histogram normalization and subthreshold-pixel elimination. **B**, Enhanced spatiotemporal Ca dynamics extracted through Penalized Matrix Decomposition (PMD) via Trefide, revealing synchronized transients, propagating waves, and localized release events within 40 x 40 pixel patches under Total Variation and Trend Filtering constraints. **C**, Residua**l** Ca activity after Principal Component Analysis (PCA) subtraction of the dominant global action-potential-driven wave, preserving intrinsic local signaling while eliminating collective propagation artifacts. **D**, Objective identi**f**ication of candidate LCRs through EVT-derived thresholding, in which Generalized Pareto Distribution (GPD) fitting across the 50th to 99th percentile range isolates high-amplitude spatiotemporal events in the extreme tail of the distribution. **E**, Composite visualization overla**y**ing EVT-identified LCRs (red) onto the original preprocessed tissue, thereby showing background Ca signals on the original data. Small red boxes in the images in panels A-D outlines the same Region of Interest inside the tissue, with spatially averaged signal shown in the respective right-hand panels as a function of time in parallel with key steps in image processing.

**Figure S2.**
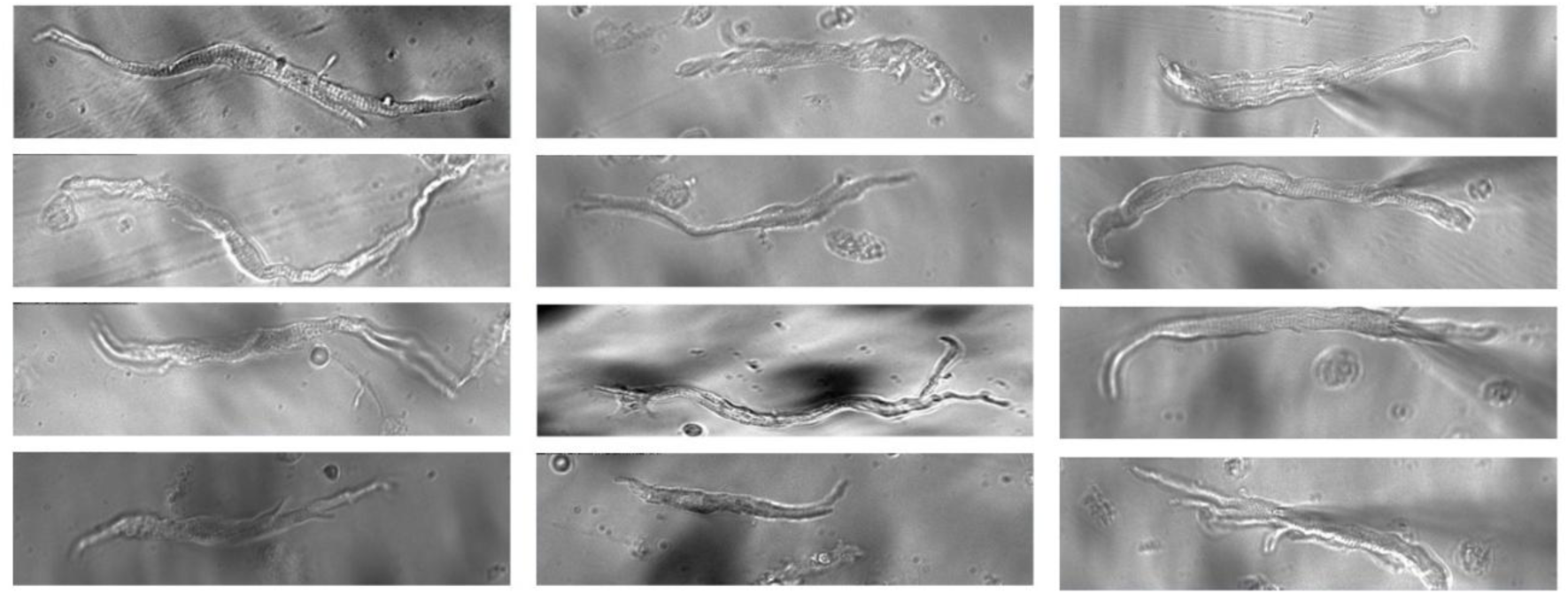
Representative examples of dormant SAN cells. The cells feature classical long curved, spindle-shaped morphology.

**Figure S3.**
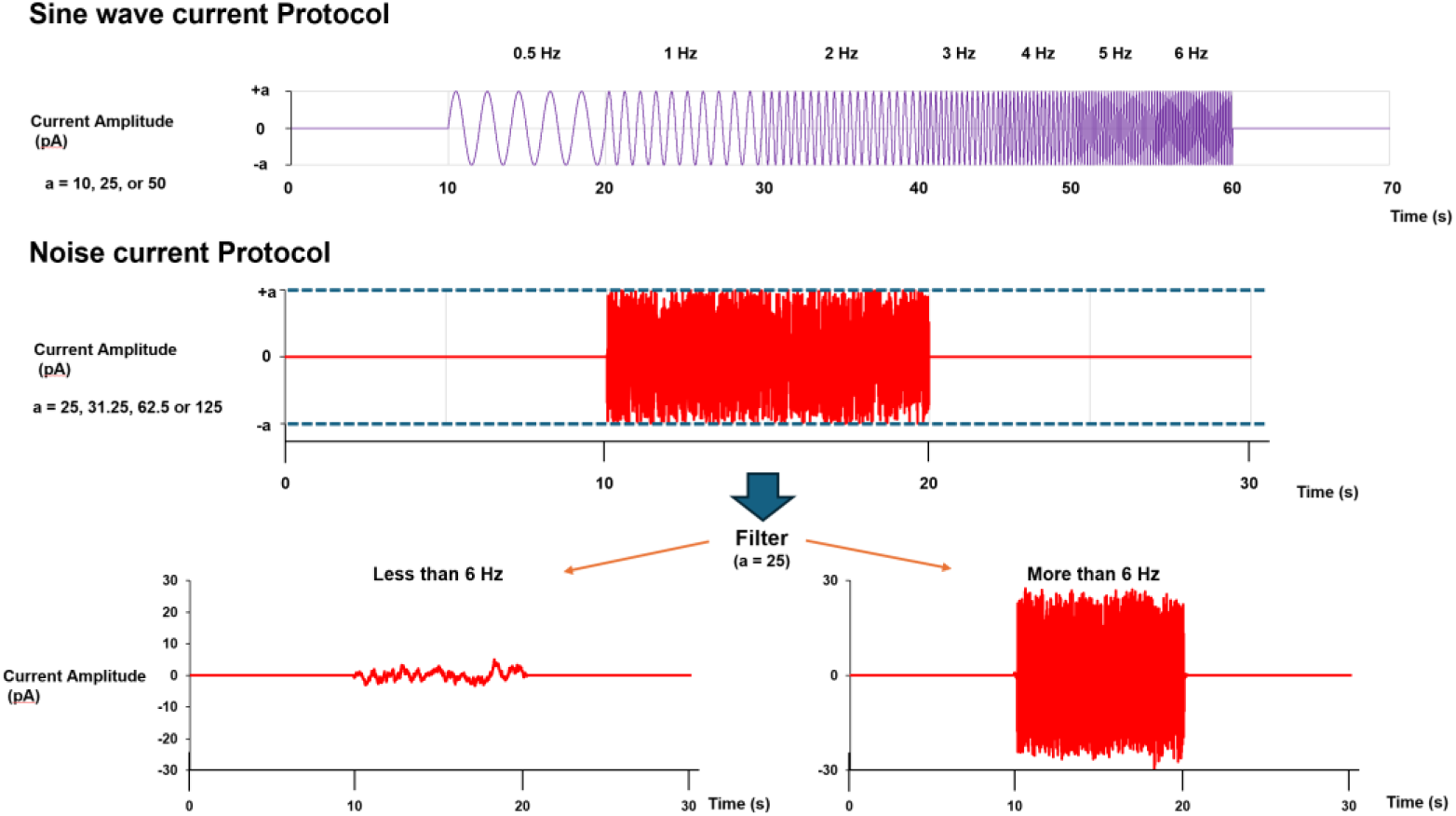
Sine wave and noise current protocols used in the study. The sine wave was applied for 50 s with sine-wave frequency increasing from 0.5 Hz to 6 Hz (upper panel). The current white noise I_noise_ was generated as a sequence of random numbers within a range of [-a, a] with an interval of 4 ms. It is important to note that while the noise amplitudes seem to be substantial, i.e. comparable with those of major currents (like I_f_ or I_CaL_), SAN cells, in fact, can process and react with one-to-one capture to only those frequency components (embedded in the white noise) which are within their resonance spectrum, i.e. below 6 Hz (the highest rate of rabbit heart, see also previous section). Thus, after a 6 Hz-low-pass filtering, the amplitude of our white noise protocol (processed by cells) decreased approximately by a factor of 5 (left bottom panel). Note, in patch-clamp experiments using Axopatch 200B patch-clamp amplifier (Molecular Devices), application of positive external currents depolarizes the cell membrane, and application of negative external currents hyperpolarizes the cell membrane.

**Figure S4.**
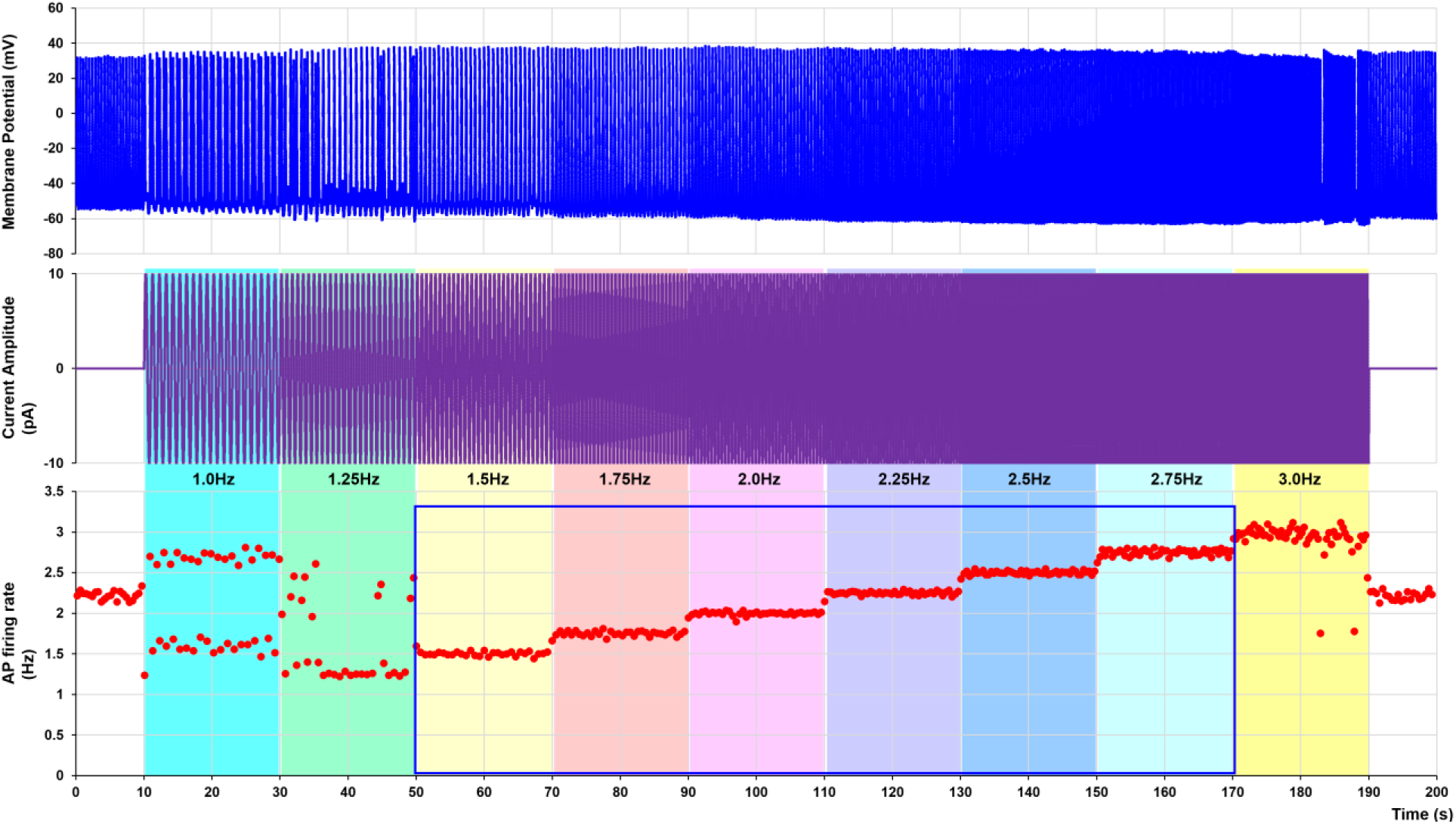
The resonance spectrum is continuous. Shown is an example of one-to-one rhythm capture in a representative cell within its resonance spectrum (from 1.5 to 2.75 Hz) to a sequence of 10 pA sine waves with a small (0.25 Hz) consecutive frequency increase.

**Figure S5.**
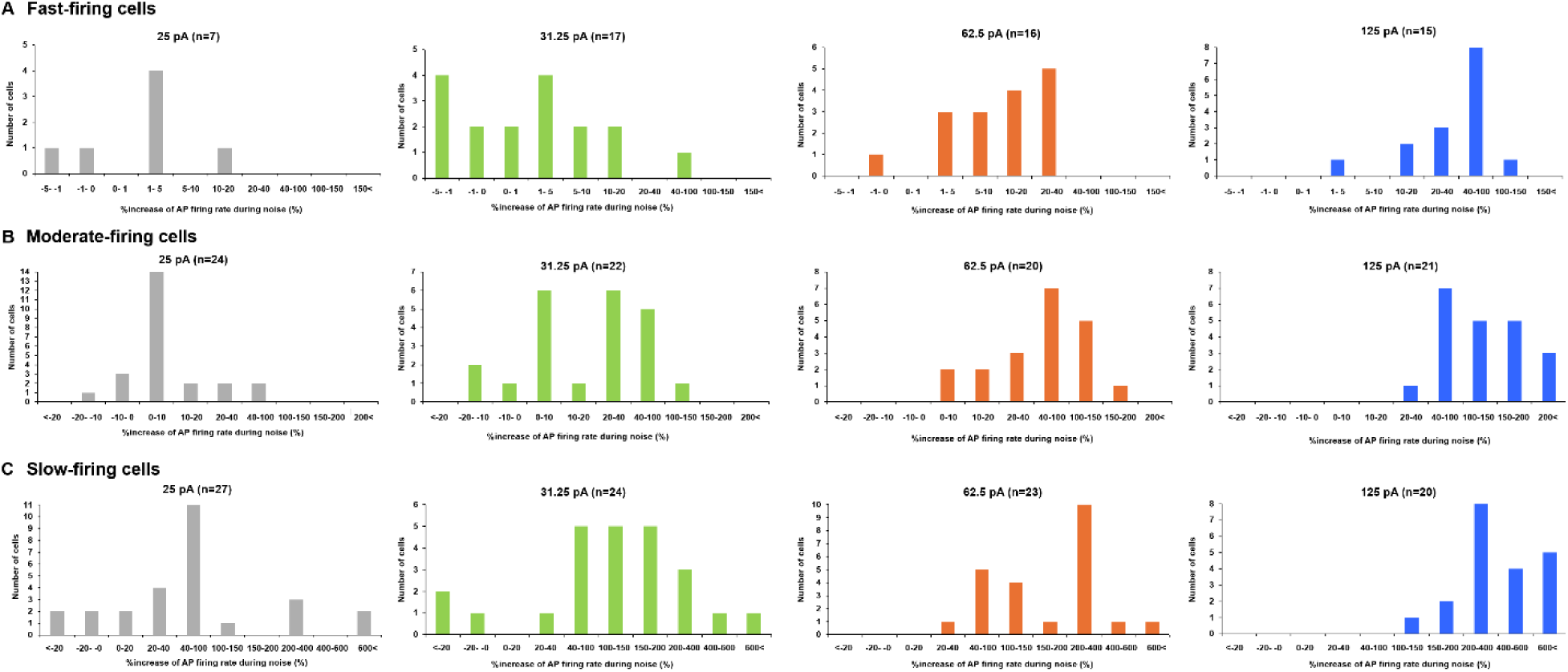
Distribution of the noise effect on the increase in average AP firing rate among all tested cells in each cell population for each noise amplitude. A, In the fast-firing group, all cells increased their average AP firing rate at 125 pA noise amplitude. Almost all cells increased their average AP firing rate at 62.5 pA noise (16/17 cells). At 31.25 pA noise, average firing rate increased in 11/17 cells. At 25 pA noise, it increased in 5/7 cells. B, In the moderate-firing group, all cells increased their average AP firing rate at 62.5 and 125 pA noise. At 25 or 31.25 pA noise, average firing rate increased in 20/24 or 19/22 cells, respectively. C, In the slow-firing group, all cells increased their average AP firing rate at 62.5 and 125 pA noise. At 25 or 31.25 pA noise, average firing rate increased in 23/27 or 21/24 cells, respectively.

**Figure S6.**
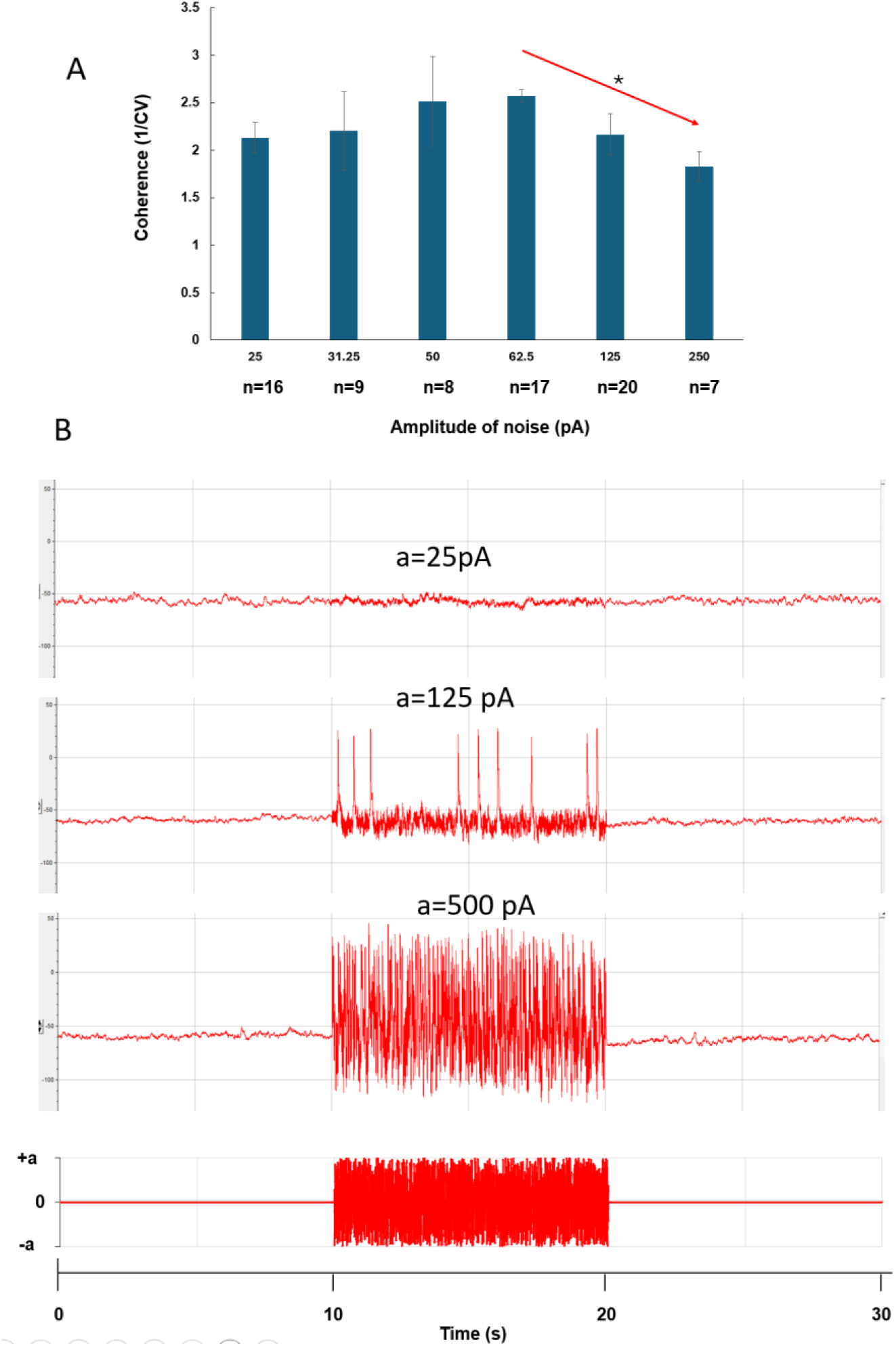
Optimal noise amplitude in stochastic resonance in a dormant cell. **A**, Temporal coherence, defined as the inverse of the coefficient of variation, revealed the classic bell-shaped profile of stochastic resonance, with firing rhythmicity being optimal at moderate noise amplitudes but low at very small and very large noise amplitudes. The Jonckheere-Terpstra trend test revealed decreasing trends in 1/CV between 62.5 pA and 250 pA (arrows; *P<0.05). **B**, Examples of original consecutive V_m_ recordings (in same cell) over a wide range of noise amplitudes, including extremely small and extremely strong noise. With extremely strong noise (bottom subpanel), V_m_ exhibited high-amplitude fluctuations deviating almost equally up and down from the baseline. AP upstrokes could not be resolved, and AP fidelity was lost.

**Figure S7.**
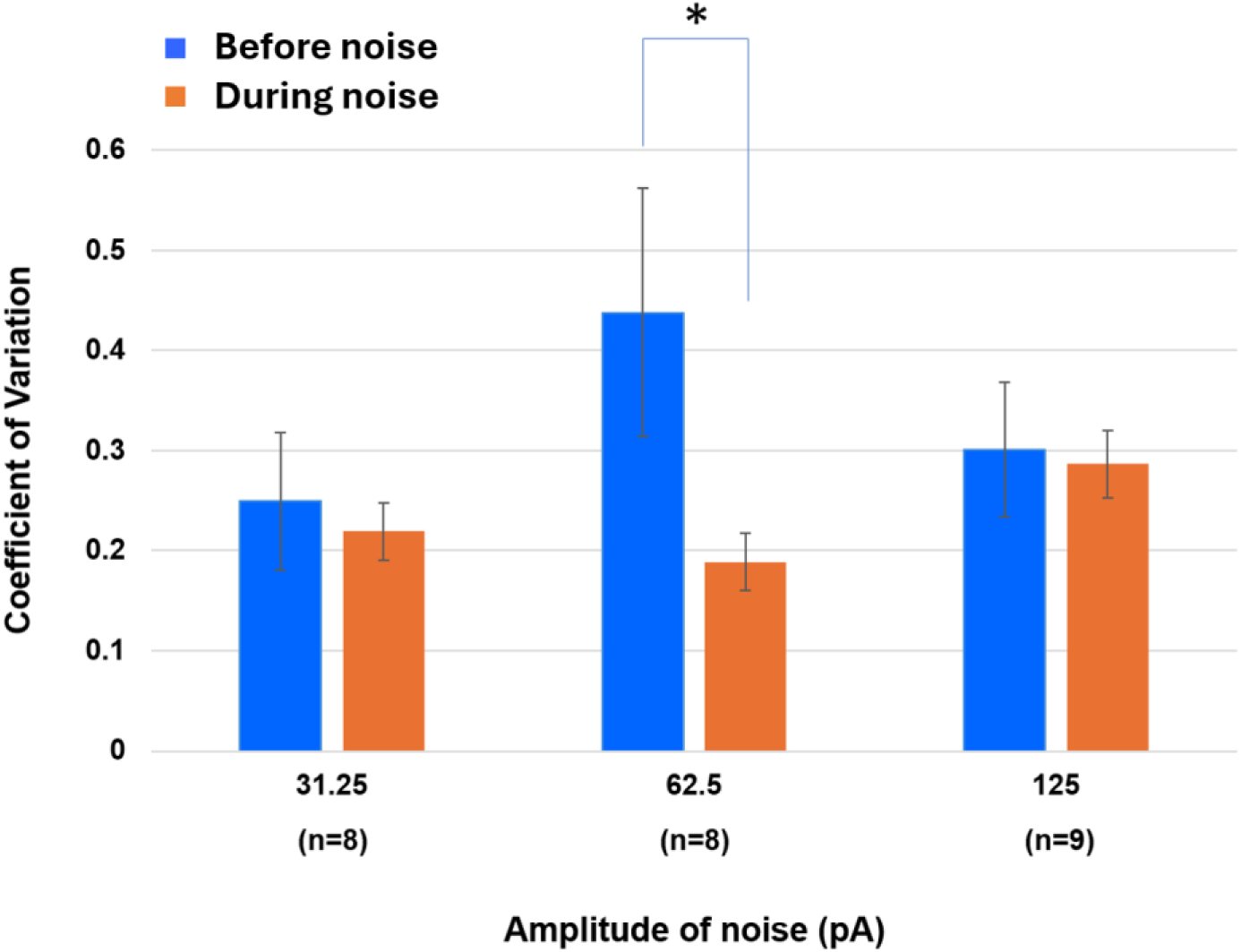
Analysis of coefficient of variation (SD/mean) of the firing frequency before and during noise of different amplitudes in cells that became slow-firing under carbachol. Substantial and statistically significant (*p < 0.05, paired t-test) decrease in CV was found only at an intermediate (optimal) noise amplitude of 62.5 pA. The decrease in CV at lower and higher noise amplitudes was small and not significant.

**Figure S8.**
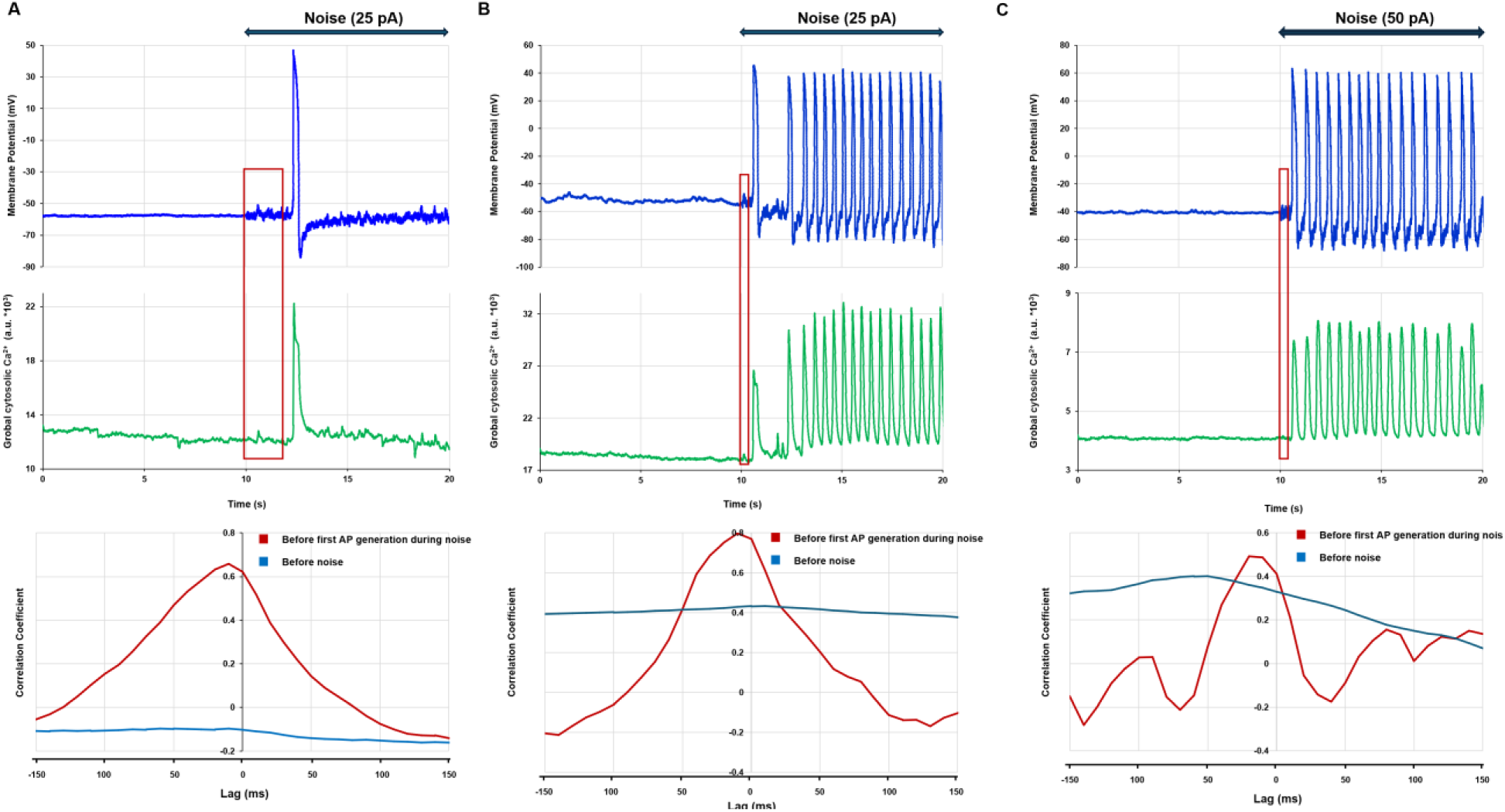
Stochastic resonance is amplified by the coupled-clock system in SAN cells. **A-C,** Representative examples of stochastic resonance in dormant cells during simultaneous perforated-patch recordings of V_m_ and Ca signals (top panels) and the corresponding cross-correlation functions of V_m_ and Ca for subthreshold signaling before and during noise application (bottom panels). The time windows selected for cross-correlation analysis (before the first AP fired) are indicated by the squares. Cross-correlation of subthreshold signals increased substantially during noise application, culminating in AP generation via stochastic resonance.

**Figure S9.**
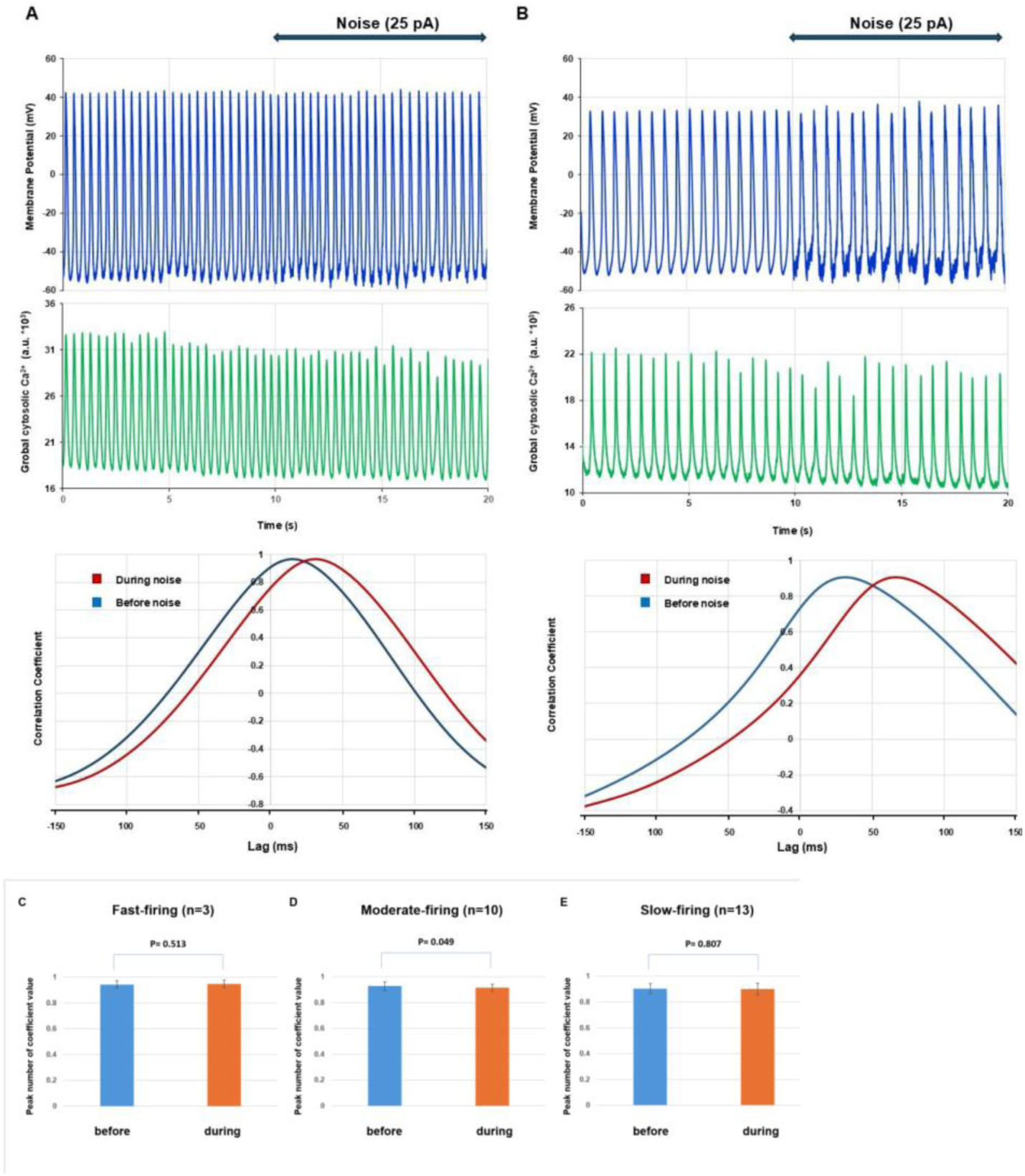
Coupling of V_m_ and Ca remains strong in the presence of noise in AP-firing-cells independent of their firing frequency. A and B: Representative examples of simultaneous perforated patch recordings of V_m_ and Ca signals (top panels) in AP firing cells in absence or presence of noise and their respective cross-correlation functions of V_m_ and Ca showing strong V_m_-Ca coupling independent of presence of noise. **C-E**: Statistical analysis showed no statistically significant change in peak value of cross-correlation in the presence of noise in all firing cell groups.

**Figure S10.**
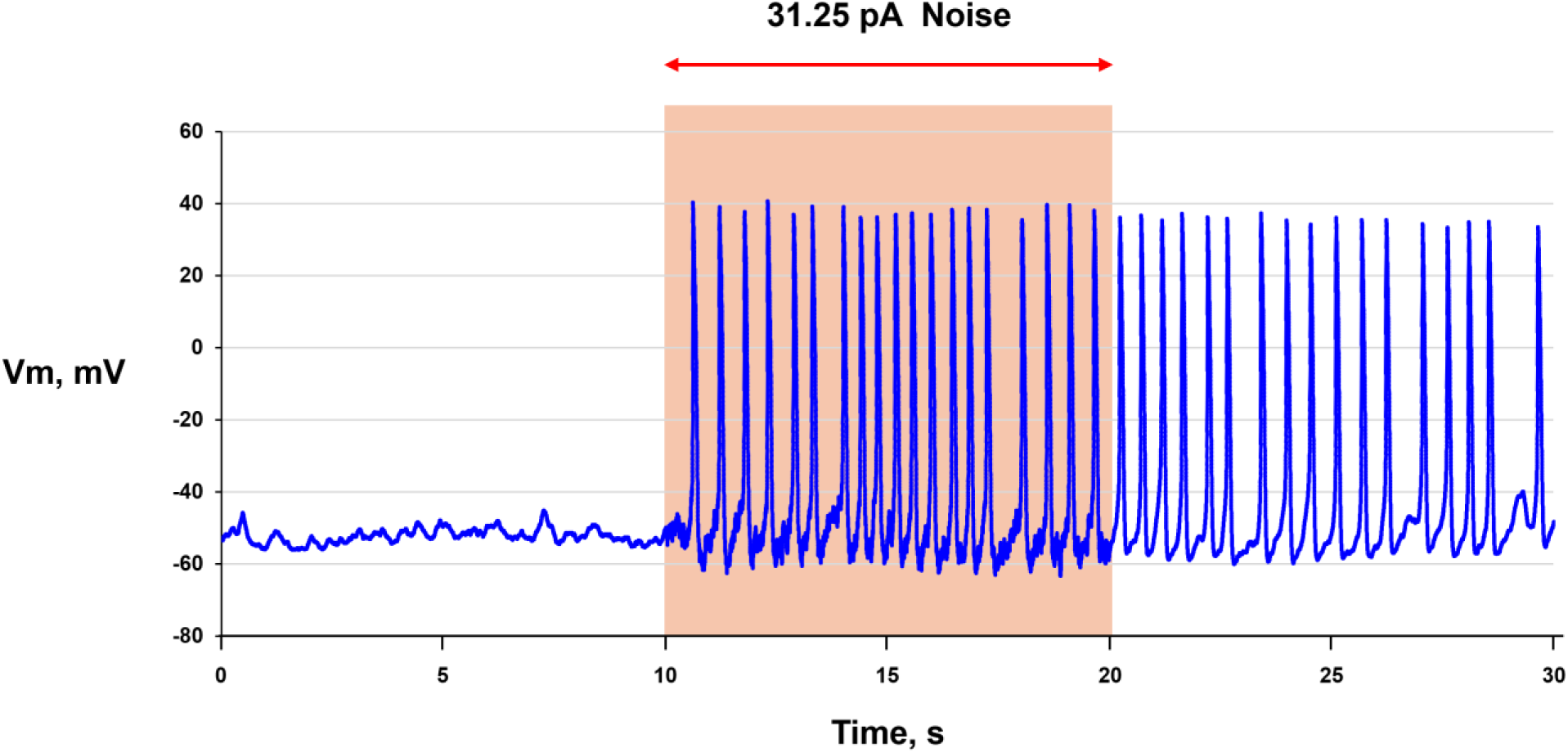
Example of a memory effect in a dormant SAN cell. Stochastic resonance awakened the cell to fire APs that continued in the absence of noise after 10 s of noise application.

**Figure S11.**
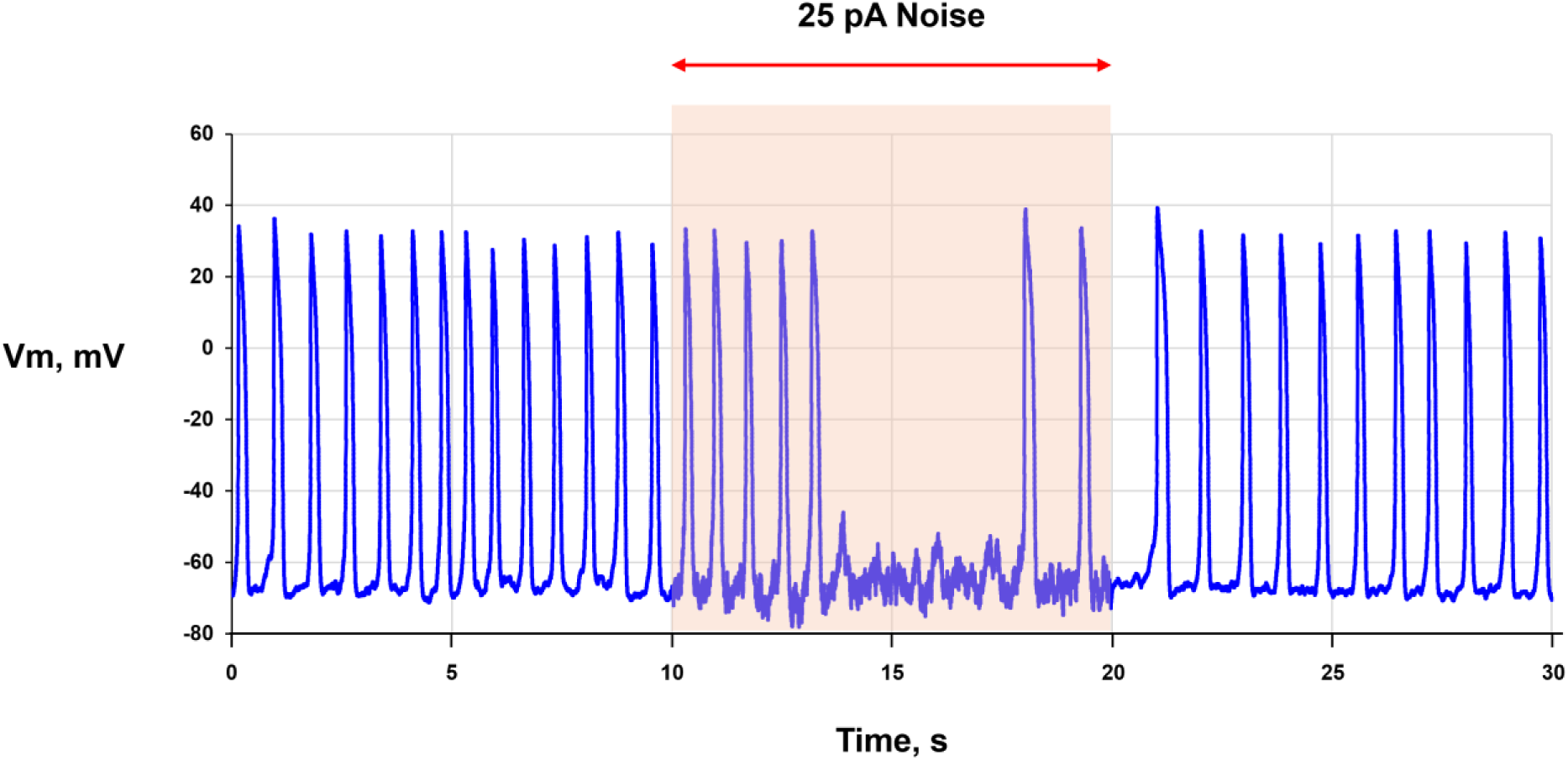
Example of inverse stochastic resonance in a SAN cell. Noise suppressed AP firing in the cell, but AP firing resumed after noise removal.

**Figure S12.**
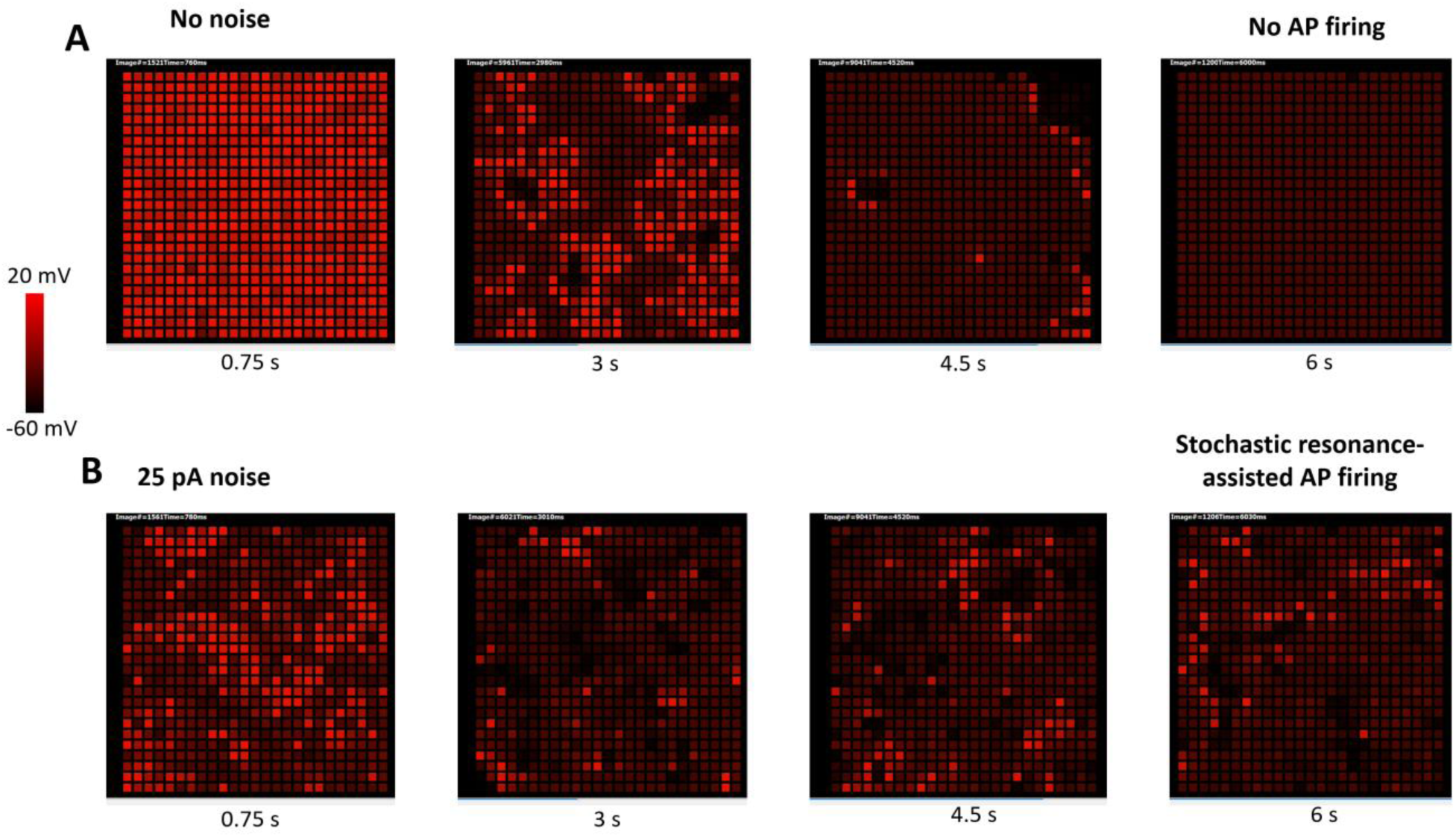
Stochastic resonance ensures fail-safe SAN operation bordering sinus arrest in our numerical simulations of SAN tissue model. Shown is an example of noise effect in a simulation of a heterogeneous SAN tissue model (25×25 cells) with P_up_ uniformly randomly distributed within the range [0, 9.5] mM/s and fixed g_CaL_=0.37 nS/pF. The model progressed into sinus arrest in the absence of noise, but continued AP firing in the presence of noise. Each subpanel shows instant distribution of V_m_ by red shades (<−60 mV as pure black to > 20 mV as pure red). See also Video S10 for V_m_ dynamics in the tissue model simulations.

**Table S1.**
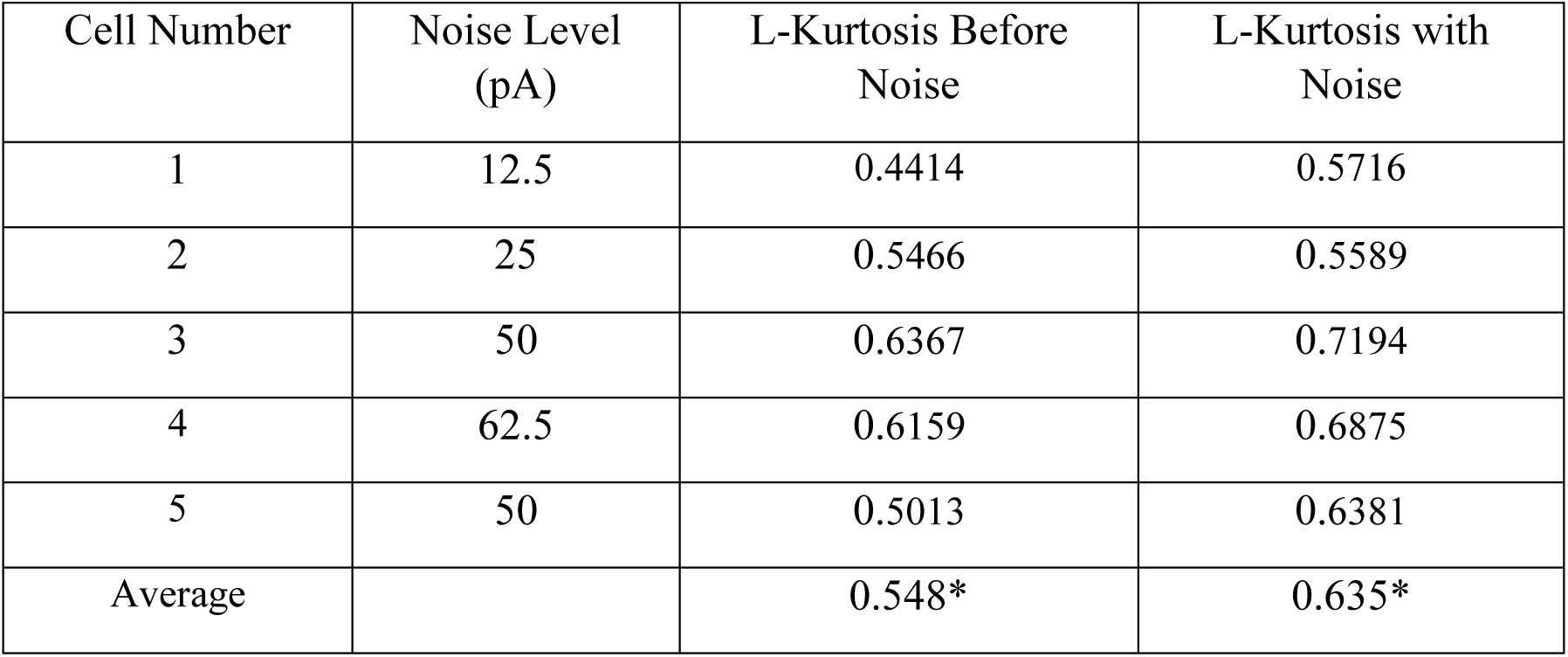
Quantitative assessment of noise-induced distributional changes using L-kurtosis, the ratio of the fourth to the second L-moment of a statistical distribution. The table shows L-kurtosis (τ4) before and during application of different noise levels to five dormant SAN cells. Higher L-kurtosis indicates a distribution with heavier tails (more outliers), whereas lower L-kurtosis indicates lighter tails (fewer outliers). All five analyzed cells showed increased L-kurtosis following noise exposure, indicating enhanced tail weight and relative enrichment of large-scale LCR events. One-tailed paired t-tests confirmed this shift toward heavier-tailed distributions (p<0.01, Cohen’s d = 1.719), supporting the conclusion that noise modulates LCR-event architecture.

| Cell Number | Noise Level (pA) | L-Kurtosis Before Noise | L-Kurtosis with Noise |
| --- | --- | --- | --- |
| 1 | 12.5 | 0.4414 | 0.5716 |
| 2 | 25 | 0.5466 | 0.5589 |
| 3 | 50 | 0.6367 | 0.7194 |
| 4 | 62.5 | 0.6159 | 0.6875 |
| 5 | 50 | 0.5013 | 0.6381 |
| Average |  | 0.548* | 0.635* |

## Legends for Videos Videos S1-S10

**Video S1. Presence of biological noise in intact SAN tissue. Top:** Low-zoom microscopic view of the entire mouse SAN preparation loaded with the Ca indicator Fluo-4, illustrating AP-induced Ca transients (APCTs) imaged by a high-speed camera. Anatomical benchmarks: crista terminalis, superior vena cava (SVC), hepatic vein (HV), and inferior vena cava (IVC). **Middle and bottom panels**: Higher-magnification views of a region of interest within the central SAN area. Local Ca release (LCR) signals detected by our novel algorithm are shown in pink.

**Video S2. Presence of biological noise in intact SAN tissue. Top:** Another high-magnification video of Ca signals in a region of interest within the central SAN area. **Bottom**: Local Ca release (LCR) signals detected by our novel algorithm are shown in pink, overlaid on the original video.

**Video S3. Examples of slow Ca waves in two SAN tissues.** Red boxes outline the locations of the slow Ca waves (low-frequency LCRs). The upper video was recorded at a sampling rate of

502.091 fps until frame #1284 (total duration 2.557 s). The lower video was recorded at 656.581 fps until frame #1736 (total duration 2.643 s).

**Video S4. Examples of multiple smaller and faster LCR signals occurring within individual cells.** Green boxes outline the locations of the LCR signals. The upper video was recorded at a sampling rate of 30 fps until frame #160 (total duration 5.3 s). The lower video was recorded at 613.333 fps until frame #1100 (total duration 1.793 s).

**Video S5. Examples of incoherent APCT firing in individual cells.** Blue boxes outline the locations of the incoherent firing signals. The upper video was recorded at a sampling rate of 644.192 fps until frame #1100 (total duration 1.707 s). The lower video was recorded at 656.581 fps until frame #1727 (total duration 2.630 s).

**Video S6. Examples of three patterns of background Ca signals detected in individual cells by confocal microsc**opy. Red circles indicate the locations of the signals. The blue box indicates a zoomed view illustrating small, frequent LCRs.

**Video S7. Example simulation of a CRU-agent model of a dormant SAN cell awakened to fire APs by application of white noise.** Spatiotemporal synchronization (black arrows) of subthreshold Ca oscillations (in green, middle panel) in the presence of noise (in grey, bottom panel). The top panel shows local Ca dynamics in red shades (10 μM = pure red; <0.15 μM = pure black) and CRUs in different functional states: ready to fire, green; refractory, blue; and releasing Ca, shades of gray reflecting junctional sarcoplasmic reticulum [Ca] dynamics (>0.3 mM = white; 0 = black). This clip is part of a longer simulation and shows 2 s before and 2 s after noise onset (outlined by a box in Figure 7A).

**Video S8. Importance of Ca-V_m_ coupling for stochastic resonance.** Representative example of stochastic resonance in a dormant cell obtained during simultaneous experimental recordings of Ca signals and V_m_, with videos of Ca signals shown before noise and during noise application (right panels). Noise increased local Ca-release activity coupled to V_m_ changes before the first AP and its associated APCT were generated (time window indicated by the yellow box in the top left panel). The increase in coupling is reflected by the higher peak of the V_m_-Ca cross-correlation function (left, bottom panel).

**Video S9. White-noise current substantially expanded the parametric space of AP firing in single-cell models (each cell in the grid represents a separate cell model).** Application of white noise in the models simulating single SAN cell function (right panels versus left panels) expanded AP firing toward where cells failed to operate without noise, both in the basal state (upper panels) and during cholinergic receptor stimulation (bottom panels). Each movie panel represents the result of numerical simulations of 5,917 (97 by 61 grid in xy) models with different coupled-clock parameters. I_CaL_ conductance (g_CaL_) was distributed evenly along the x-axis from 0.28 to 0.52 nS/pF in 0.0025-nS/pF increments. The sarcoplasmic reticulum Ca pumping rate (P_up_) was distributed evenly along the y-axis from 0 to 12 mM/s in 0.2-mM/s increments. White noise of 25 pA amplitude was added to each individual (not connected) SAN cell model in the grid. V_m_ dynamics in each model are coded by red shades from black (<-60 mV) to pure red (>20 mV). Image number and simulation time are shown in the upper left corner of each video panel. See also Figure 8A-D.

**Video S10. Example of the effect of noise in simulations of the SAN tissue model**. The SAN model progressed into sinus arrest after a few synchronized cycles in the absence of noise (left panel), whereas in the presence of noise the SAN model continued AP firing (right panel). Each cell in the square grid of the SAN model is represented by a box colored in red shades corresponding to V_m_ from <-60 mV (pure black) to >20 mV (pure red).

## Glossary of Key Terms

**Sinoatrial node (SAN)**: A small region of specialized cardiac tissue in the right atrium that spontaneously generates APs to pace the heartbeat.

**Dormant cell**: A SAN cell that generates subthreshold oscillations but does not produce APs under basal conditions; can be awakened by adrenergic stimulation or noise.

**Stochastic resonance**: A phenomenon in which the addition of an optimal level of noise enables a subthreshold signal to reach detection threshold, improving system output.

**Inverse stochastic resonance**: A phenomenon in which noise suppresses rather than enhances oscillatory activity, observed here in some fast-firing cells.

**Subthreshold signal**: A signal whose amplitude is insufficient to independently trigger an AP.

**Resonance spectrum**: The range of external signal frequencies to which a given SAN cell responds with one-to-one AP capture; equivalent to the cell’s functional frequency processing range.

**Coupled-clock system**: The interaction between membrane ion channel oscillations (membrane clock) and intracellular Ca cycling (Ca clock) that together drive rhythmic AP generation in SAN cells.

**Local Ca release (LCR)**: A spontaneous, localized intracellular Ca release event generated by ryanodine receptor channels in the sarcoplasmic reticulum during diastolic depolarization.

**White noise:** A random signal with equal power at all frequencies, used here to simulate the broadband biological noise present in SAN tissue.

**Coefficient of variation (CV)**: The ratio of standard deviation to mean of AP firing frequency, used as a measure of firing rhythmicity; lower CV indicates more regular firing.

**AP-induced Ca transients (APCTs)**: periodic synchronized flashes of synchronized Ca releases throughout the SAN tissue induced by respective action potential firing via classical Ca-induced Ca-release (CICR) mechanism.

**Ca-induced Ca-release (CICR)**: Ca release that is triggered by Ca entered via L-type Ca channels (classical CICR) or by a local neighboring Ca release, resulting in propagating Ca release in the form of Ca wave that, in turn, could be short abrupted wave (wavelet or LCR) or full-cell wave (global Ca wave).

**References: see reference list in the main text**

